# High resolution structures of Myosin-IC reveal a unique actin-binding orientation, ADP release pathway, and power stroke trajectory

**DOI:** 10.1101/2025.01.10.632429

**Authors:** Sai Shashank Chavali, Peter J. Carman, Henry Shuman, E. Michael Ostap, Charles V. Sindelar

## Abstract

Myosin-IC (myo1c) is a class-I myosin that supports transport and remodeling of the plasma membrane and membrane-bound vesicles. Like other members of the myosin family, its biochemical kinetics are altered in response to changes in mechanical loads that resist the power stroke. However, myo1c is unique in that the primary force-sensitive kinetic transition is the isomerization that follows ATP binding, not ADP release as in other slow myosins. Myo1c also powers actin gliding along curved paths, propelling actin filaments in leftward circles. To understand the origins of this unique force-sensing and motile behavior, we solved actin-bound myo1c cryo-EM structures in the presence and absence of ADP. Our structures reveal that in contrast with other myosins, the myo1c lever arm swing is skewed, partly due to a different actin interface that reorients the motor domain on actin. The structures also reveal unique nucleotide-dependent behavior of both the nucleotide pocket as well as an element called the N-terminal extension. We incorporate these observations into a model that explains why force primarily regulates ATP binding in myo1c, rather than ADP release as in other myosins. Integrating our cryo-EM data with available crystallography structures allows the modeling of full-length myo1c during force generation, supplying insights into its role in membrane remodeling. These results highlight how relatively minor sequence differences in members of the myosin superfamily can significantly alter power stroke geometry and force-sensing properties, with important implications for biological function.

**Significance Statement:** Myosin-IC (myo1c) uses an ATP-driven ‘power-stroke’ to support slow intracellular membrane and vesicle transport. We used cryo-electron microscopy to understand adaptations of myo1c to perform its unique roles. We discovered an altered interface between myo1c and actin compared with the closely related myo1b, which repositions the motor domain and alters the trajectory of its lever arm swing compared to other myosins. This explains why myo1c propels actin filaments in a leftward circular path. We also discovered a unique role in force sensing for a structural element called the N-terminal extension and built a full-length atomic model for the myo1c power-stroke. Our findings highlight how myosins can tune their power-stroke geometries and force-sensing properties to adapt to diverse cellular functions.

## Introduction

Members of the myosin family of cytoskeletal motors function in a wide range of cell processes that include muscle contraction, organelle transport, cell adhesion, signal transduction, and cell division. The motor domain of each myosin paralog evolved mechanochemical properties suited to carry out these diverse functions (1, 2). Notably, the ATPase activities and motility rates of myosin paralogs can differ by three orders of magnitude (2, 3), with strikingly different sensitivities to mechanical load resulting in motors with varied force-velocity relationships and power outputs (1). Determining how sequences within the highly conserved motor domains lead to such diverse mechanochemistry has been a challenge to the field.

Myosin-Is are a family of single-headed, membrane-associated motors that function in cellular processes related to membrane morphology and trafficking, tension sensing, regulation of actin dynamics, and nuclear transcription (4, 5). Like the myosin superfamily overall, there is substantial mechanochemical diversity within the myosin-I family (5). For example, two widely expressed vertebrate paralogs, myo1b and myo1c, have similar slow biochemical kinetics (6–9). However, optical trapping studies show that the actin detachment rate of myo1b is slowed ∼100-fold by mechanical loads < 2 pN (10, 11), while myo1c is relatively insensitive to loads in this range (8). This dissimilarity is due to differences in the sensitivity of ADP release to mechanical loads. Functionally, myo1b has mechanochemistry suited to force-sensitive actin anchoring, while myo1c is suited to generating power under loads (1). An additional interesting difference is that myo1c is able to turn actin filaments in leftward circles in gliding assays (12), which is a property that may be related to establishment of cell chirality (13).

In previous work, we determined the high-resolution structures of rigor and ADP-bound, tension-sensitive states of actin-bound myo1b from rat (14). Biophysical studies based on these structures revealed a role for the N-terminus of the protein in communicating the presence of mechanical loads to the nucleotide binding site, which affect ADP release (14–16). We termed this region the N-terminal extension (NTE) and showed that it interacts with the N-terminal subdomain of the motor, the lever arm helix, and the converter domain. Its sequence is variable among myosin-Is, and similar regions exist in other myosin paralogs (16–18); however, it is not present in myosin-V (19). Although the importance of the NTE has been established for modulating the rate of ADP release (15, 16), it is not clear how sequence differences in the motor lead to altered force sensitivity.

The ability of myosins to adjust their kinetic properties in response to force is important for tuning force-velocity relationships in muscle (20–23), facilitating processive stepping of transport motors (24), and maintaining tension of membranes (10). Interestingly, the biochemical step of myo1c that is most sensitive to mechanical loads is different from other myosins that have been studied, including myo1b, myo5, myo6, smooth muscle myosin (MYH11), and cardiac muscle myosin (MYH7). While other characterized myosins respond to force by slowing ADP release, myo1c responds by slowing the isomerization that follows ATP binding (8). The origin of this fundamental difference in behavior of myo1c has remained obscure. In part, this is because the structural origins of ADP force-sensitivity also remain incompletely understood.

In this work, we report the high-resolution structures of the ADP-bound and rigor (AM) states of actin-bound myo1c (actomyo1c) expressed without its membrane-binding tail domain (residues 1-767; see Methods) and we compare the structures to actin-bound myo1b. We discovered that myo1c binds to actin in a unique orientation that produces a ‘skewed’ power stroke with respect to the actin filament, and that this effect is enhanced by an inherent skew of the myo1c power stroke itself, compared with myo1b. The skewed power stroke may explain the motor’s ability to turn actin filaments in gliding assays (12). Moreover, we find differences from myo1b in the structural relationship between the nucleotide binding site and position of the lever arm helix that provide a rationale for differences in force sensitivity. Finally, our new structures allow us to model the full working stroke of the full length myo1c molecule, providing insights into function of the native molecule. These results provide for a more complete understanding of the coupling of the ATPase cycle and lever arm position of myosins.

## Results

### Two-step isomerization visualized during the myo1c ADP lever arm swing

We generated complexes of a mouse myo1c construct containing three IQ motifs and calmodulin bound to actin filaments and solved atomic-resolution structures both in the absence and presence of 1 mM MgADP (Fig. 1). The resolution of the myo1c motor domain (Fourier shell correlation, 0.143) was estimated as 2.7 Å for the rigor state (AM) and 2.8 Å for the ADP-bound state (AM.ADP; Fig. S1). This resolution allowed modeling of the protein chains throughout the actin, motor, and first IQ motif (Fig. 1). The resolutions of the lever arm helix and calmodulins beyond the first IQ motif were substantially lower than those above, so we did not include these regions in the final structures.

**Fig. 1.**
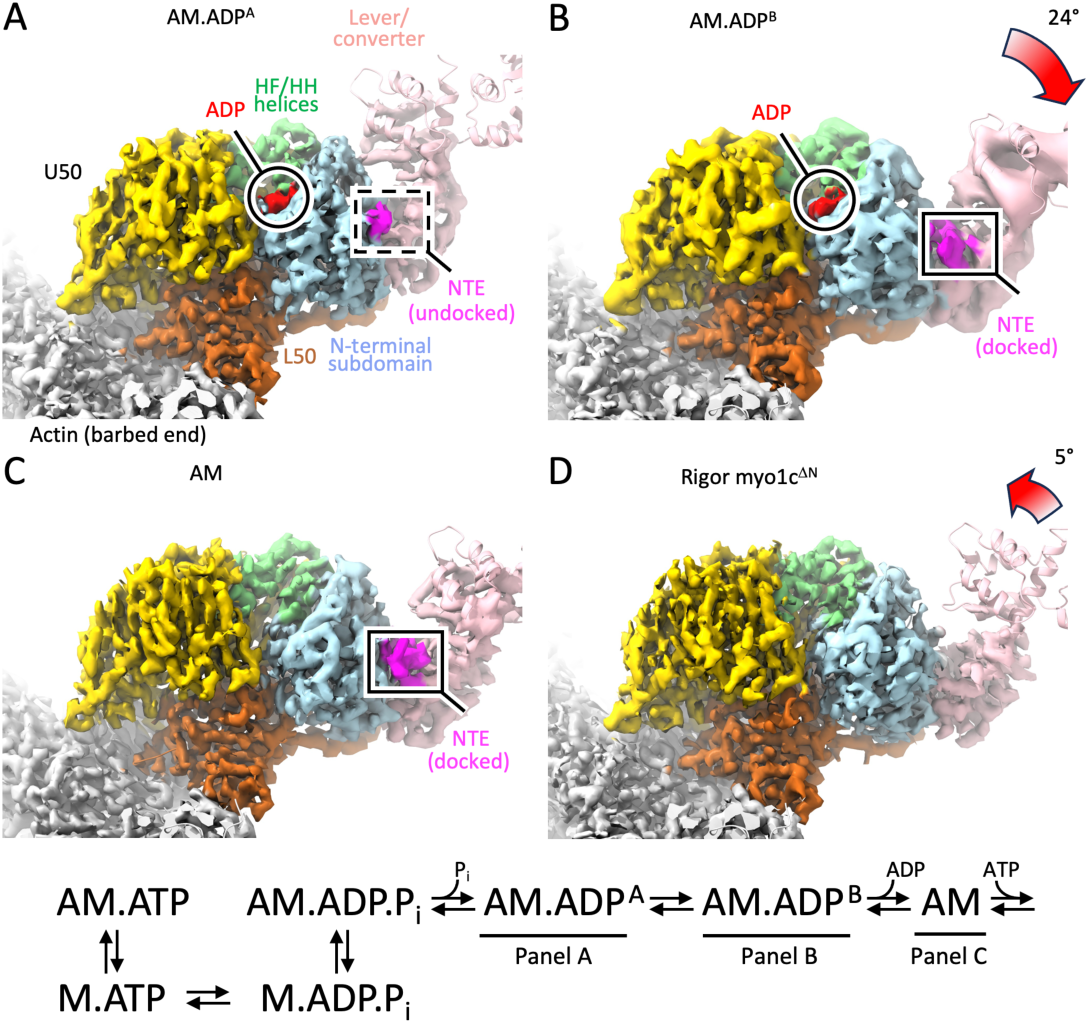
ADP-dependent lever swing of myo1c revealed by near-atomic cryo-EM structures. (*A*) View of the 2.8 Å resolution structure of actin-bound ADP myo1c, in the AM.ADP^A^ state. The actin barbed-end points out of the page, slightly obliquely, toward the reader. Myosin subdomains are color coded as: (yellow) upper-50 kDa domain; (caramel) lower 50-kDa domain; (light blue) N-terminal subdomain; (light green) HF and HH helices; (light pink) lever arm plus converter; (red) ADP; (magenta) N-terminal extension. Only a few C-terminal residues of the NTE are ordered (dashed box). (*B*) Cryo-EM structure of the myo1c AM.ADP^B^ state with identical coloring and view as AM.ADP^A^. The lever arm swings 24° relative to AM.ADP^A^, and density for docked NTE bridges between the lever and the N-terminal subdomain. (*C*) AM state of myo1c cryo-EM structure, showing a lever position very similar to AM.ADP^B^. (*D*) AM myo1c^ΔN^ cryo-EM structure, showing a lever that reorients ∼7° back towards the AM.ADP^A^ position compared with wild-type. Density maps are shown as colored isosurfaces superposed on the molecular models, which are rendered with ribbon cartoons. To improve visibility of lever features, the isosurface contour threshold level is locally decreased in all four panels (see Fig. S1). The actomyosin ATPase scheme is shown with correlated structural states identified by figure panel.

The structures reveal a tilting of the lever arm going from a principal ADP-bound state conformation (AM.ADP^A^) to the AM state (Fig. 1; Fig. S2 - S4) as seen previously at low resolution (9). Similar to what was observed for actin-bound myo1b, but unlike other myosins that have been studied (14), 3D classification revealed a second ADP-bound population (AM.ADP^B^) whose lever arm has repositioned to a rigor-like orientation (Fig. 1B; Fig. S3). Classification analysis indicates this AM.ADP^B^ population may comprise ∼5% of the total particles ( Fig. S4). The myo1c lever tilts 24° from AM.ADP^A^ to AM.ADP^B^ and < 2° from AM.ADP^B^ to AM (Fig. 1). Corresponding myo1b lever arm rotations were 25° and 5°, respectively (14); thus, we propose that, similar to myo1b, the three myo1c conformational states give a mechanistic succession of states proceeding from AM.ADP^A^ to AM.ADP^B^ to AM (14).

Despite these similarities, however, myo1c AM.ADP^B^ and AM conformations are not the same as in myo1b. In myo1c, conformational changes within the motor domain that accompany ADP release appear largely complete in the AM.ADP^B^ structure, and the AM structure differs little from AM.ADP^B^ (0.7 Å backbone RMSD; Table S1). In contrast, the myo1b nucleotide cleft opens only partially in AM.ADP^B^, and further opening of the nucleotide cleft with ADP release results in an AM conformation distinct from AM.ADP^B^ (1.2 Å RMSD; Table S2). As shown below, differing behavior of the myo1c and myo1b motor domains during ADP release is linked to significant functional differences that are observed between these two motors.

### An off-axis myo1c ADP lever arm swing

Cryo-EM models reveal that the actin-bound myo1c lever arm swing upon ADP release is skewed 32° from the long axis of the actin filament, while the myo1b ADP lever arm swing is much more parallel (7° skew) (Fig. 2). When modeled in the context of an actin gliding assay, skewing of the myo1c ADP swing is predicted to push the leading tip of an actin filament to the left (Fig. S5-S7). These observations provide a plausible mechanism by which myo1c promotes left-handed circular, rather than straight, gliding of actin filaments in *in vitro* motility assays (12, 13).

**Fig. 2.**
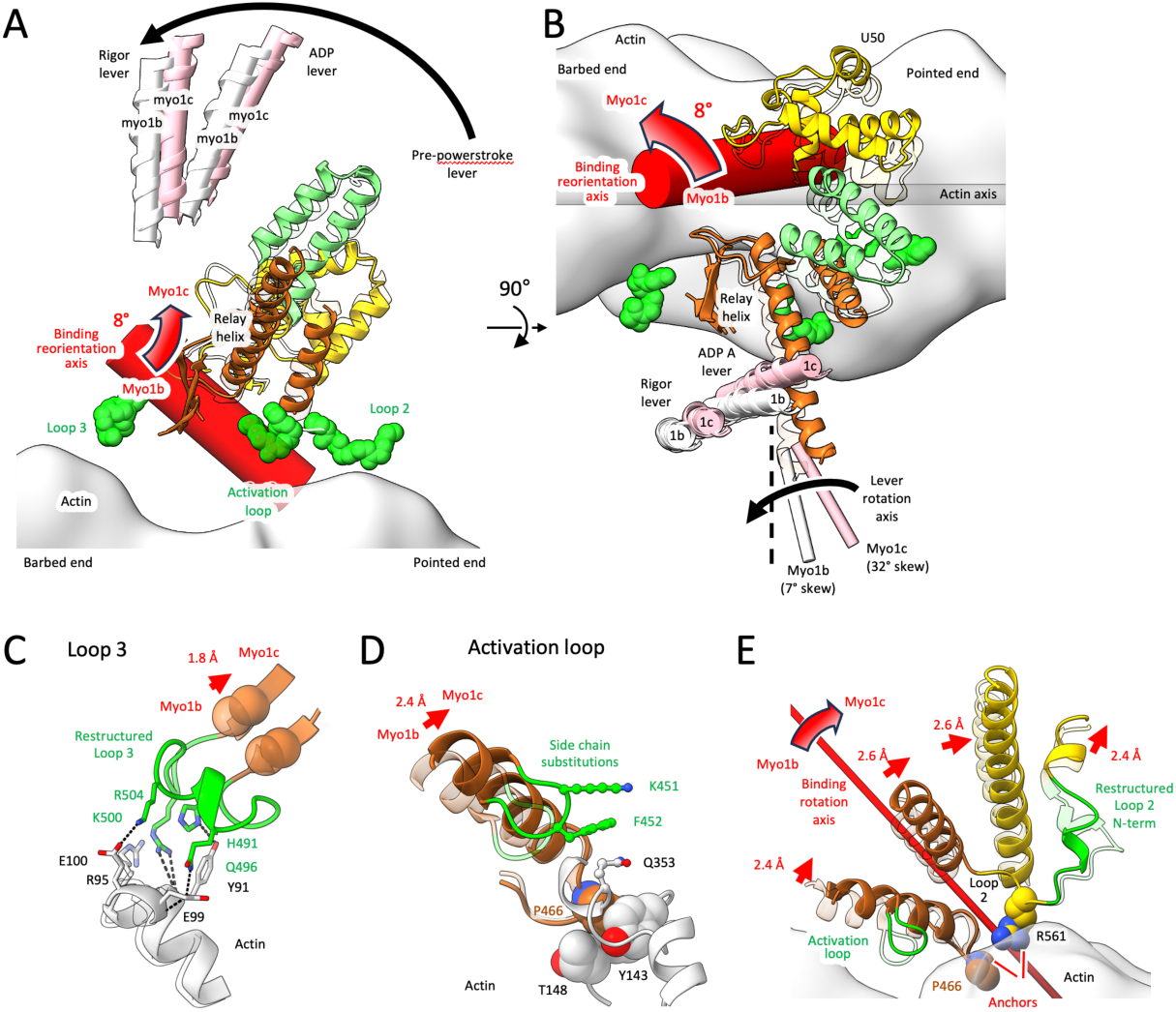
Structural origin of a skewed lever swing in myo1c. See also Figs S6 and S7 and movies S0 and S1. (*A*) Side view of acto-myo1c (actin axis is horizontal) showing eight-degree rotation of the myo1c motor domain on actin compared with myo1b. Myo1b (14) (transparent ribbons) and myo1c cryo-EM structures (colored ribbons) are aligned by their central three actin subunits. Starting from this alignment, a thick red cylinder depicts the binding reorientation axis required for least-squares superposition of rigor myo1b upper and lower 50-kDa domains onto myo1c; the axes for ADP-state superpositions are similar (not shown). The reorientation tilts the lever arm helix (pink ribbons with cylinders running through them, and white ribbons and cylinders for myo1c and myo1b, respectively) towards the actin pointed-end. Three myosin loops whose actin contacts are linked to the binding orientation change are depicted as green van der Waals spheres. Full details of the binding reorientation axis calculation are given in the Movie S1 caption. (B), Orthogonal view of (A) revealing that the ADP lever arm swing of myo1c is more skewed (larger off-axis component) than myo1b. Lever rotation axes defining the hinge points of the myo1b and myo1c ADP lever swings are depicted as thin cylinders, colored white and pink, respectively. Full details of lever arm rotation axis estimation are given in Fig. S7. For reference, the black vertical dashed line denotes a lever rotation axis with no off-axis component (*C*) Restructuring of myo1c loop-3 that accommodate its repositioning on the actin surface compared with myo1b (see Fig. S6A). In this and the following panels, structure alignments are the same as (*A*)–(*B*) and the viewing angles are close to *(A*), but individually adjusted for best viewing of selected elements. (*D*) Side chain substitutions in the myo1c activation loop that accommodate its repositioning on the actin surface compared with myo1b (see Fig. S6B). (E) Restructuring of myo1c loop-2 accommodates the orientation change (see Fig. S6C–D and Movie S2). Conserved side chains that serve as anchor pivots for the myo1b to myo1c re-orientation are represented using van der Waals spheres. Note that, different to the Fig. 2A-B, the binding reorientation axis is rendered in this panel as a thin rather than thick cylinder

More detailed comparison of the myo1c and myo1b ADP lever arm swings reveals two distinct factors that both contribute to the off-axis lever arm swing in myo1c: the binding orientation of the motor domain on actin, and changes in the lever arm swing geometry inherent to the motor domain itself. The binding orientation, or perch, of the myo1c motor on actin is different from myo1b (Fig. 2A–B). The myo1c perch is defined by distinctive interactions at the actin-binding interface (Fig. 2; Movie S2; see below). The myo1c actin-binding region in our structures (upper- and lower-50 kDa domains) remain mostly static in our structures, evidenced by minimal movements of the two domains relative to each other (no more than 1– 3° rotation; see Fig. 3A-B). The lower 50-kDa domain can therefore be used as a measure of the motor orientation change with respect to actin.

**Fig. 3.**
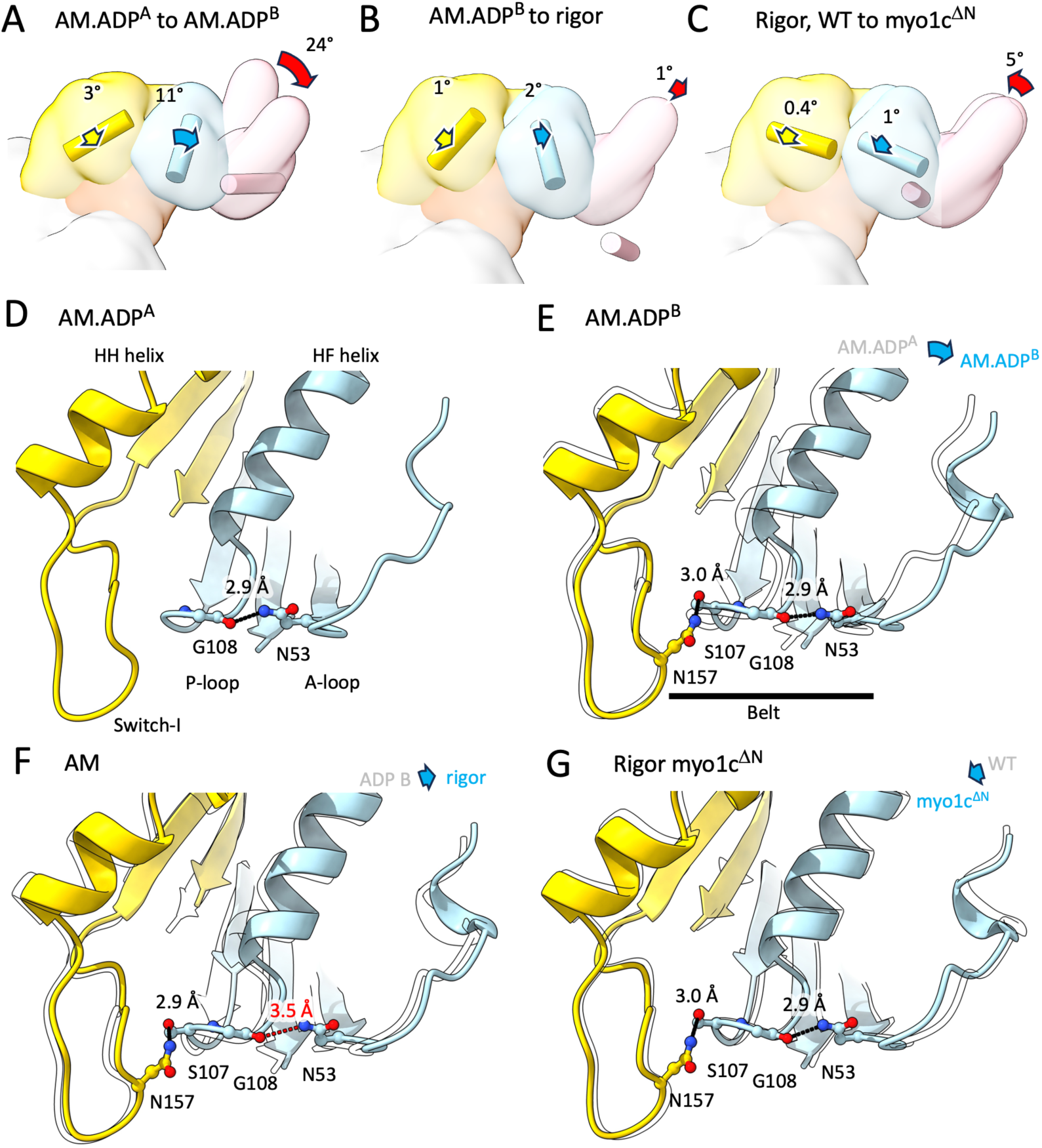
Nucleotide-dependent domain movements related to ADP release from actin-bound myo1c. (*A*) Subdomain rotations comparing our AM.ADP^A^ and AM.ADP^B^ structures, from a reference alignment where lower 50 kDa subdomains are superimposed. The myo1c lever/converter and other domains are represented as isosurfaces of 16 Å resolution synthetic density maps generated from our atomic models of AM.ADP^A^ and AM myo1c; domains are colored as in Fig. 1. Colored cylinders depict the rotation axis required to superimpose respective myo1b and myo1c subdomains with respect to the reference alignment. Colored arrows denote the rotation direction. Subdomains rotate very little except for the (pink) lever/converter where a significant lever swing is evident. The lever rotation axis (pink colored cylinder) is positioned at the structural hinge location as in Fig. 2A-B. The remaining rotation axes are positioned at the respective subunit centers of mass. (*B-C*) Analysis as in (*A*) showing that nucleotide-binding subdomain rotations going from AM.ADP^B^ to AM are subtle and are partially reversed in the AM myo1c^ΔN^ structure. (*D-G*) Details of the nucleotide pocket for the four myo1c structures. Atomic models are rendered as ribbon cartoons with stick models for selected residues; reference ribbon structures corresponding the comparisons in (*A-C*) are overlayed as transparent silhouettes. Strained H-bond between the P-loop and A-loop is colored red in (*F*). Subdomain rotations and shifts were estimated using published python scripts (51) within UCSF ChimeraX (57).

In our myo1c structures the actin-binding region rotates 6–8° counterclockwise compared to myo1b when looking down on the motor perpendicular to the filament axis (Fig. 2B; Movie S1). This orientation change redirects the ADP lever arm swing of myo1c, making it ∼4° less parallel to the filament long axis than if it assumed the myo1b perch orientation (Fig. S7B and Movie S1, compare green to pink cylinders; see also Fig. S7C). We expect the myo1c primary lever arm swing from the pre-power stroke state represented by (PDB:4BYF) to AM.ADP^A^ would be similarly affected. Strikingly, while perch differences are evident in other reported actomyosin structures, the motor domains tend to reorient in a manner that would not be expected to skew the lever arm swing significantly compared with myo1b (Fig. S8). We note that a low-resolution structure of *Acanthamoeba* myosin-IB was also reported to have a unique binding orientation on actin (25), but it remains unclear what affect this would have on the lever arm swing trajectory.

Skew of the myo1c AM lever arm swing caused by the unique perch orientation is greatly enhanced by unique lever arm geometries inherent to the motor domain itself. We quantified this effect by computing the lever arm rotation axis that would result if the myo1c motor domain assumed the same perch orientation on actin as myo1b (Fig. S7B and Movie S1, green cylinders). The resulting skew of this ‘myo1b-oriented’ myo1c ADP lever swing, 28°, is much larger that of the myo1b ADP lever swing (7°) (Fig. S7B and Movie S1, white cylinders). The total myo1c ADP lever swing skew (32°; Fig. S7B, pink cylinders) is even larger due to the 4° perch difference.

Comparing lever arm orientations when motor domains are aligned reveals that the skewed myo1c lever swing is mainly due to a difference with the AM lever arm position. Lever orientations of aligned myo1b and myo1c AM.ADP^A^ motor domains are within ∼2–3° of one another (Fig. S7C, compare middle black and green data points; Tables S5–S6). However, lever orientations of aligned AM structures differ more substantially between the two isoforms, resulting in a significant off-axis component (azimuthal angle change) of the ADP lever arm swing for myo1c but not myo1b (11° and 3°, respectively; Fig. S7C, right side and Tables S5–S6). This inherent difference between myo1b and myo1c AM lever arm positions may originate from different structural environments of the lever in this state; in particular, in myo1b loop-5 maintains contact with the lever in all three structural states (AM.ADP^A^, AM.ADP^B^ and AM), while this contact is lost in the myo1c AM.ADP^B^ and AM structures (Movie S1; also see below). In summary, compared with myo1b, myo1c skews its ADP swing off-axis by concomitantly changing its perch on actin and adjusting the AM lever position.

### Structural determinants of the myo1c perch orientation on actin

The cryo-EM density of the myo1c actin-binding site reveals well-defined side chains and other features consistent with the reported resolution of 2.7 - 3.0 Å (Fig. S2) and is essentially indistinguishable in the AM.ADP^A^, AM.ADP^B^, and AM states. Thus, we used the highest resolution structure (AM) to identify changes at the myo1c actin binding site compared with myo1b that drive reorientation of the perch (Fig. 2B-E; Movie S2). Overall, myo1c actin-binding loops on the periphery of the interface principally drive the reorientation, as was observed in structural studies of PfMyoA (26).

Sequence and structure differences at three distinct sites of the myo1c actin interface drive the unique orientation: Loop-2 (K553 – T564) in the upper 50-kDa domain, and loop-3 (T490 – E506) and the activation loop (E450–G454) in the lower 50-kDa domain. Reorientation occurs about a fixed pivot point on actin, where conserved P466 in the lower 50-kDa domain rests in a hydrophobic pocket (Fig. 2D, E). Notably, the cardiomyopathy loop and loop-4 have unique structures, but are positioned similarly to other characterized myosins (27).

Loop-2 (K553 - T564) connects the upper and lower 50-kDa domains while interacting with actin where it plays a role in reorienting myo1c on actin. While loop-2 is largely disordered in the Myo1c.ADP.VO_4_ (PDB: 4BYF; (28)) crystal structure, it is well ordered and resolved in the three actomyo1c states. The loop-2 C-terminus is fixed on actin near the P466 pivot in both myo1c and myo1b, supported by hydrogen bonds of R561 with actin S344 and S348 (Fig. S6C, D). However, a helix-loop-helix motif immediately N-terminal to loop-2 operates as a spacer that lengthens the loop by ∼2.5 Å in myo1c, holding the upper 50 kDa domain farther away from the actin surface than myo1b (Fig. S6C; Movie S2). A major determinant of the spacer restructuring in myo1c is the loss of two prolines from the myo1b spacer sequence (myo1b P555, P559). These sites surround a glutamic acid (myo1b E556; myo1c E555) that makes a conserved salt bridge with an arginine (myo1b R359; myo1c R353) in a neighboring α-helix of the upper 50-kDa domain. These differences and other sequence changes, including D552, K553, and S554 in myo1c that orients and stabilizes an α-helical turn S555-S558) drive the spacer lengthening. The spacer function we identify for loop-2 in myo1b and myo1c may be related to another posited role of loop-2, which is to mediate the initial interaction between some myosins and actin (27, 29), thereby regulating actin-activated ATPase activity.

Loop-3 (T490-E506) in the lower 50-kDa domain is the other major actin interface that accommodates the myo1c binding reorientation. Myo1c loop-3 shifts ∼2 Å away from the actin surface compared with myo1b and other myosins (Fig. 2C; Fig. S6A; Movie S2). Substitution of extended hydrophilic side chains (Q496, R499, K500 and R504) in place of shorter counterparts in myo1b (N496, D502, T503 and H507, respectively) supports this shift, as well as restructuring of the actin-proximal part of this loop into an α-helical turn to shift R499 and K500 backbone positions closer to the actin surface in myo1c (Fig. 2C; Fig. S6A). When myo1c binds to actin, actin Y91 is flipped 120° compared to when myo1b binds. This reorientation allows Y91 to fill a vacant space in the myo1c structure that would otherwise be occupied by the more closely positioned loop-3 of myo1b. As a result, despite being farther away from actin than in myo1b, myo1c loop-3 maintains a similar number of actin contacts, with both electrostatic and hydrophobic character. Interestingly, the actin-detached myo1c.ADP.VO4 structure (PDB: 4BYF; (28)) shows a similar loop conformation to the cryo-EM structures, suggesting that the loop’s shape is maintained by its internal structure.

The myo1c activation loop (E450-G454) is also shifted on the actin surface compared with myo1b (Fig. 2D). The activation loop is a short region between the HQ and HR helices in the lower 50-kDa region of myosin (30). Upon actin binding, it is thought to accelerate the movement of the relay helix, which stimulates ATPase activity (31). F452 of myo1c forms a hydrophobic interaction with Q353 and a possible backbone interaction with S350 of actin. In contrast, the residue corresponding to F452 in the myo1b activation loop (T448) does not interact with actin, and other actin interactions of the loop are minimal, limited to a single backbone hydrogen bond of N447 with actin S350 (Fig. S6B). Although not resolved, K453 likely interacts with the N-terminal acidic residues (E4) of actin.

The cardiomyopathy loop (T319-P333) in the upper 50-kDa region also plays a significant role in the myo1c binding orientation change. However, unlike loop-3 and the activation loop, the cardiomyopathy loop is positioned similarly on actin as other characterized myosins (27) and maintains a similar (although not identical) structure. Instead, to compensate for the different motor domain orientation of myo1c, the cardiomyopathy loop swivels ∼5° on the upper 50-kDa subdomain compared with myo1b. The new cardiomyopathy loop orientation relative to the motor is supported by orientation changes of the short helices at the base of loop-2 due to hydrophobic repacking. In particular the short helix in myo1c 535-539 is shifted such that it follows the cardiomyopathy orientation change. The cardiomyopathy loop contains residues that form hydrophobic interactions observed in other myosins (I323, A325, L330). The side chains of S332 and R321 of myosin form a unique charge cluster with side chain of E334 of actin, and E328 of myo1c interacts with K336 and Y337 of actin. The K336 and Y337 interactions are intriguing, because these residues are involved in the “flattening” of the actin subunit as it polymerizes to form F-actin. Given the proposal that some myosins affect actin nucleation and polymerization (32), the effect of this interaction on actin dynamics should be explored.

Other actin-interacting regions of myo1c contribute less to the binding orientation change. The helix-loop-helix motif adjacent to loop-3 is situated very close to the binding orientation pivot axis (not shown), so that actin interactions are very similar to other myosins, and the interacting residues are more conserved. The helix-loop-helix motif stabilizes the DNase binding loop (D-loop) of actin, as found for other myosins (27). This includes the side chain of myosin’s E476 with K50 of actin. There is a complex network of interactions that both stabilize the helix-loop-helix structure and interact with two adjacent actin subunits, including E461, E462, and K477 of myosin with T351 and G46 of actin. L472, L475, and L478 of myosin form a hydrophobic patch that positions loop-3.

Loop-4 (N281-E297) in the upper 50-kDa subdomain is differently composed than in myo1b and is located farther from the actin surface. In contrast to hydrophobic actin contacts made by myo1b loop-4, E288 backbone and D289 side chain atoms from myo1c loop-4 possibly interact electrostatically with K328 of actin; however, these residues are poorly resolved. The sequence and orientation of this loop likely explain the competition between myo1c and tropomyosin for binding to actin, as shown in *in vitro* motility and biochemical experiments (33, 34).

### Opening of the active site accommodates ADP release

The overall behavior of myo1c during ADP release, as captured by our structures, generally parallels myo1b; however, differences in the details correlate with highly significant functional differences between the two motors. Both motors make a large-angle lever swing (∼25°) during ADP release (9, 35), accompanied by a significant rotation (∼10°) of the N-terminal domain to open the nucleotide pocket, and both motors dock their NTE segments into a cavity between the motor domain and the lever in the AM conformation. Moreover, both motors exhibit small sub-populations of the ADP state (AM.ADP^B^) where the lever has swung to an angle that approaches the AM position. This behavior differs from other myosins, where multiple ADP conformations have not been reported. However, details of nucleotide pocket opening and its coordination with NTE docking and lever arm tilting during ADP release differ substantially between myo1c and myo1b. These structural relationships are likely crucial for setting force sensitivity.

The global conformational changes in myo1c that accompany active site opening during ADP release are largely completed in the AM.ADP^B^ state. Subdomain motions of the three states can be described by approximate rigid body rotation of the converter/lever arm helix and the N-terminal subdomain (G12-R97 and K592-G626) in relation to the remainder of the motor domain. The myo1c AM.ADP^A^ to AM.ADP^B^ transition is accompanied by a 10° rotation of the N-terminal subdomain against the upper 50-kDa subdomain, reflecting opening of the active site; very little rotation of the subdomains is then observed in the AM.ADP^B^ to AM transition (< 1°). A related observation is that, while density corresponding to Mg^2+^ is pronounced in the myo1c AM.ADP^A^ nucleotide pocket (Fig. S3A), it is weak or absent in the AM.ADP^B^ structure (Fig. S3B), presumably because the nucleotide pocket opens to a rigor-like conformation that disrupts the Mg^2+^ binding site (19). In contrast, the rotations of the myo1b N-terminal subdomain are equally divided between the AM.ADP^A^ to AM.ADP^B^ transition (5°) and the AM.ADP^B^ to AM transition (5°), reflecting incremental openings of the active site that occur in each step (see also Table S2). Reflecting this difference, Mg^2+^ evidently remains able to bind in myo1b AM.ADP^B^, prior to the cleft opening more fully in AM to release Mg^2+^ and the nucleotide (14).

The opening of the myo1c active site during ADP release occurs through the movement of the HF helix away from the HH helix, increasing the distance between switch-1 and the P-loop (Fig. 3). At their ends where HF and HH are connected by loop-1 (T125 -A128), these helices in myo1c remain closer to each other than seen in myo1b. The other ends of the helices make a minor “chopstick” movement going from AM.ADP^A^ to AM.ADP^B^, such that the P-loop and switch-1 loop translate away from each other by ∼1.2 Å. This movement is very different from the movement of the HF helix observed in myo1b and myo5 (14, 36), where there is a significant axial helix-helix sliding component.

During the transition from AM.ADP^A^ to AM.ADP^B^, the P-loop and the C-terminally connected HF helix move laterally ∼2 Å away from switch-1 and its N-terminally connected HH helix. During this transition, hydrogen bonds are formed between N157 in switch-1 and S107 in the P-loop. Despite high conservation of these residues, this interaction was not seen in myo1b or myo5, but was reported in myo15 (14, 32, 36). It is notable that this P-loop switch-1 interaction combines with a highly conserved H-bond between neighboring G108 of the P-loop and N53 of the A-loop (S51 - R63; (36)) in the N-terminal subdomain, making an H-bond interaction “belt” in AM.ADP^B^ that holds the active site in a conformation permissible for nucleotide release and binding (Fig. 3E).

The conformational transition from myo1c AM.ADP^B^ to AM is subtle compared with myo1b. This is likely due to the P-loop to switch-1 H-bond constraining the P-loop movement. As a result, the HF helix/P-loop element fails to make a ‘piston-like’ axial movement away from switch-2 seen in the corresponding myo1b transition (14). However, the myo1c transition from AM.ADP^B^ to AM features a ‘breathing’ movement of the upper 50 kDa domain and N-terminal subdomains (Fig. 3B-C, Fig. S9, Movie S2) accompanied by perturbation of the H-bond between the P-loop and N53 (2.9 to 3.6 Å; Fig. 3D, E). This ‘breathing’ movement thus correlates with opening of the AM active site in a manner that would likely be less amenable to strong nucleotide binding.

### Y75 and E76 in loop-5 have key roles in the transition of the lever from AM.ADP^A^ to AM.ADP^B^

Similar to the changes in the active site, tilting of the myo1c lever and accompanying movements in the N-terminal subdomain are largely complete in the transition from AM.ADP^A^ to AM.ADP^B^. There is little change in the positions of the lever, NTE, and N-terminal subdomain in the subsequent transition to AM (Fig.’s 1, 3; Movie S3).

Two critical residues (Y75, E76) in loop-5 (R70 - H80) are involved in the transition of the lever from myo1c AM.ADP^A^ to the conformation seen in AM.ADP^B^ and AM (Fig. 4). In the AM.ADP^A^ state these residues form a bridge between the motor domain and the lever (Fig. 4C-E). Y75 in loop-5 is found in a hydrophobic pocket with L688 and F689 of the converter, which we term the ‘L5–lever bridge,’ while E76 forms a cluster of interactions with residues in the converter and lever arm (R639, Y640, E692).

**Fig. 4.**
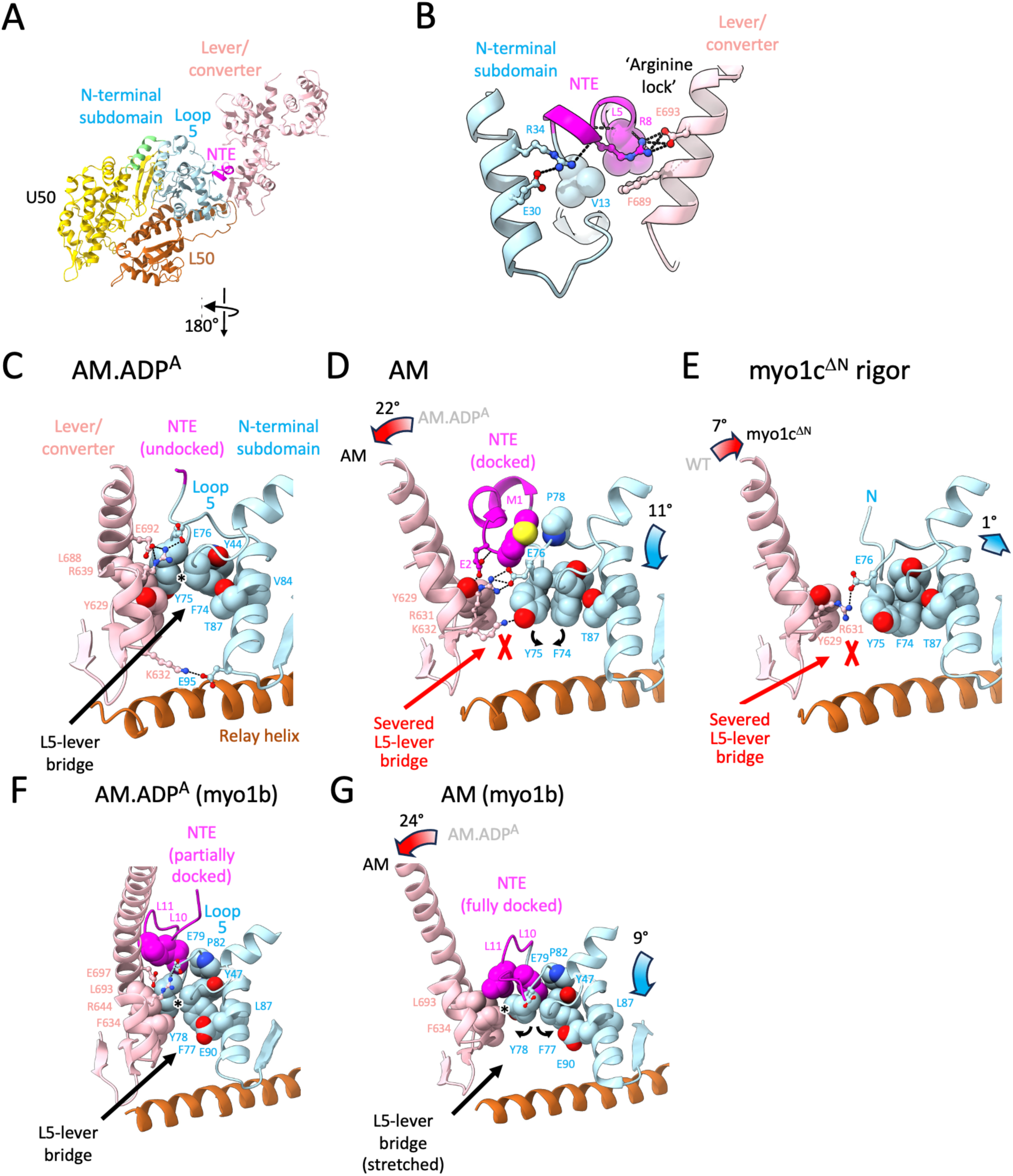
Conformational changes in the myo1c and myo1b lever arm and N-terminal extension associated with ADP release and force-sensitivity. (*A*) Overview of the myo1c structure, shown in the AM state with a docked N -terminal extension; viewing angle and coloring are the same as in Figs. 1 and 3. (*B*) Close-up of the docked N-terminal extension in (*A*), highlighting electrostatic interactions (ball-and-stick-rendered side chains) unique to myo1c that lock the lever orientation with respect to the N-terminal subdomain, including a cation-pi stacking interaction between R8 in the NTE and F689 in the lever. A hydrophobically interacting side chain pair is depicted using VDW spheres. (*C - E*) Angle-dependent changes in lever-motor domain interactions for different myo1c conformational substates. View is of the same region as in (*B*), but from the opposite side (180° rotation around y). Residues depicted as VDW spheres are involved in a hydrophobic contact bridge (’L5–lever bridge’) between the lever and loop-5, which severs upon lever rotation from the (*C*) AM.ADP^A^ to (*D*) AM states. Curved arrows indicate interaction changes of residue pair F74-Y75 in loop-5. Cryo-EM density in this region of the AM structure is essentially indistinguishable from the AM.ADP^B^ structure (Fig. S10). (*E*) Myo1c^ΔN^ AM state. (*F - G*), Corresponding region in myo1b AM.ADP^A^ and AM structures (14), showing that the L5–lever bridge distorts but does not break during ADP release.

Rotation of the N-terminal subdomain from AM.ADP^A^ to the AM.ADP^B^/AM states concomitantly opens the nucleotide cleft and moves L5 away from the lever. This movement disrupts the L5–lever bridge by pulling Y75 out of the converter hydrophobic pocket and breaking a cluster of interactions between E76 and the lever (Fig. 4C–D). Disruption of this bridge allows the lever to tilt, completing the ADP swing. While L5 is evidently more mobile in our AM.ADP^B^/AM cryo-EM maps and density for these side chains is weak or absent, modeling indicates that Y75 repacks with neighboring F74 in loop-5, and E76 forms a new cluster of interactions with the converter (Y629 and R631) and the docked NTE (E2, see below), supporting the AM.ADP^B^/AM lever position (Fig. 4E).

The equivalent of Y75 is conserved in myo1b (Y78) but does not interact with the neighboring F74 (F77) in AM.ADP^B^ or AM. Rather, it adopts a different rotamer that maintains hydrophobic contact with the lever following the ADP lever arm swing, and blocks the lever from tilting back. Thus, in contrast to myo1c, the L5–lever bridge in myo1b does not break as the lever swings during the transition to AM.ADP^B^ and AM states. Instead, the bridge stretches and maintains contact with the lever by swiveling the aromatic residue pair (F77, Y78) away from each other in AM.ADP^B^ and AM conformations (Fig. 4G).

The differing behavior of the L5–lever bridge in the two motors is supported by side chain substitutions in myo1c that alter motor domain contacts made by the conserved phenylalanine. In myo1b the F77 side chain conformation is kept in a similar conformation throughout ADP release by hydrophobic contacts with an adjacent helix (side chains L87 and E90). E90 also sterically blocks F77 from pivoting towards Y78 (Fig. 4F-G). In myo1c, L87 and E90 are replaced by smaller side chains (V84 and T87) that are too short to contact F77, thus enabling F77 to pivot towards Y78 in the AM.ADP^B^ and AM conformations, breaking the hydrophobic bridge as ADP is released (Fig. 4D-E). Additionally, the E2 interaction cluster (Fig. 4D) is not observed in myo1b, as the myo1b NTE blocks the ability of E76 (E79 in myo1b) to interact with the conserved residues in the converter. The L5–lever bridge appears to be a critical determinant of force-sensitivity; in particular, failure of myo1c to maintain this bridge throughout the ADP lever arm swing likely disables ADP force-sensitivity (see Discussion).

### Myo1c NTE has a unique structure that stabilizes the lever arm position

The NTE is a crucial participant in the AM.ADP to AM transition, as it markedly accelerates the rate of ADP release and mechanically stabilizes the AM state under load (15, 16). Accompanying the opening of the active site and tilting of the N-terminal subdomain in the AM.ADP^A^ to AM.ADP^B^ transition, we observe NTE docking in a pocket formed by the N-terminal subdomain, converter, and lever arm (Fig. 4).

The docked NTE locks the myo1c lever into place in AM.ADP^B^ and AM through a network of interactions that are dominated by cation-pi, salt bridge, and H-bond interactions. R8 has a prominent role in this network that we call an ‘arginine lock,’ forming an apparent cation-pi bond with F689, and simultaneously forming a salt bridge with D693 to stabilize the lever arm position (Fig. 4B). R8 also forms a backbone hydrogen bond interaction with L5 of the NTE. The arginine-lock is not present in myo1b, where NTE docking is dominated by hydrophobic interactions. M1 of the docked myo1c NTE makes hydrophobic contacts with V77 and P78 of loop-5.

### Deletion of the myo1c NTE disrupts the AM lever position and alters the nucleotide pocket

Deleting the myo1c N-terminus (E2-V10; myo1c^ΔN^) substantially slows the rate of ADP release, eliminates the ADP-release-associated tilting of the lever, and increases the rate of ATP binding to rigor actomyo1c (16). To investigate the structural origins of this behavior, we solved the cryo-EM structure of rigor (AM), actin-bound myo1c^ΔN^ (16) at a resolution of ∼2.7 Å (hereafter, referred to as ‘myo1c^ΔN^ rigor’). This structure reveals that in the absence of an intact NTE, the lever moves toward the motor domain, intruding into the space that in the wild-type AM structure is occupied by the docked NTE. Loss of M1 results in disorder of part of loop-5 (72-75). Our structure model also indicates that E76 reorients to occupy the space vacated by M1 (Fig. 4E; Fig. S10), although the E76 side chain itself is not directly visualized presumably due to its net negative charge. Moving closer to the motor domain leads the myo1c^ΔN^ lever to make a new putative interaction with loop-5 (E76-R639) in addition to the E76-R631 interaction observed in the wild-type AM.

Notably, these changes in lever and loop-5 conformations in the myo1c^ΔN^ rigor structure compared with wild-type rigor (AM), correlate with changes at the nucleotide pocket that appear to regulate nucleotide affinity. The A-loop, which connects to the C-terminus of loop-5, moves with the N-terminal subdomain domain (∼1°) back toward the P-loop to allow formation of the H-bond between N53 and the P-loop H-bond, thus restoring the H-bond interaction belt that connects the switch-1, P-loop, A-loop. If this belt facilitates tight nucleotide binding to the P-loop, loss of coupling to the lever/loop-5 in the myo1c^ΔN^ mutant may explain the enhanced rate of ATP binding to myo1c^ΔN^ (see below).

### Modeling the power stroke of full length Myo1c

We built a full length myo1c (FL-myo1c; Fig. 5) molecule allowing us to model the trajectory of the power stroke using the previously determined crystal structure of the myo1c tail domain that includes the lever arm helix and three bound calmodulins (PDB: 4R8G; (37)). The region from the end of the tail to the motor domain does not contain hinges or regions of substantial flexibility, suggesting that the myosin can be considered rigid between the actin- and membrane-binding sites (Fig. 5).

**Fig. 5.**
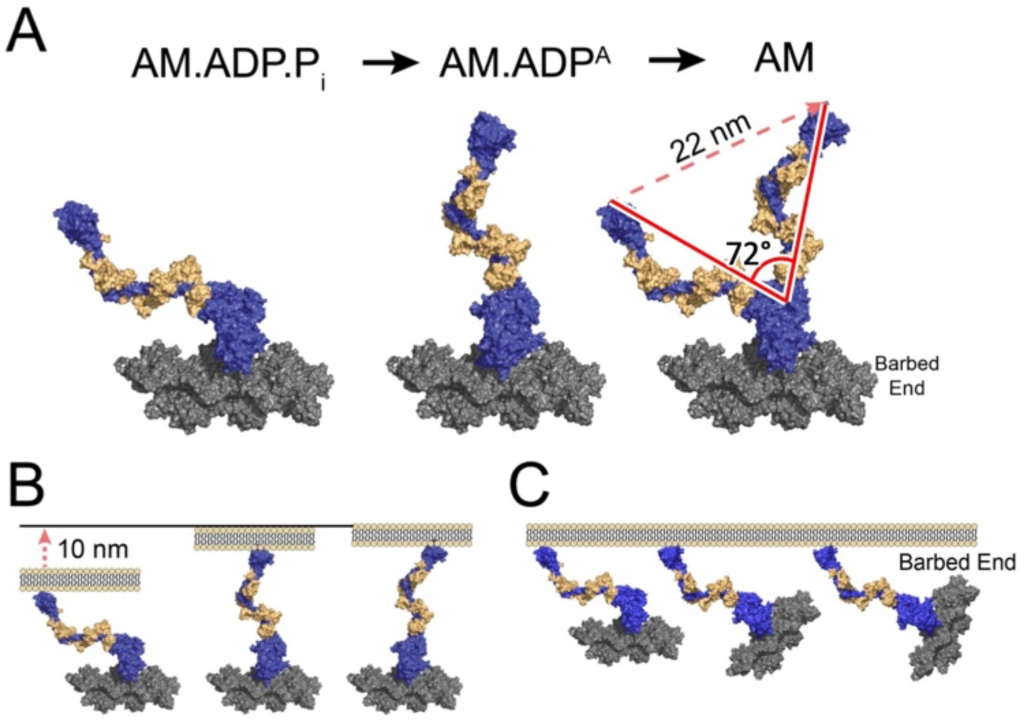
Full-length myo1c model complexed with actin suggests roles in membrane remodeling. (*A*) VDW sphere representation depicting the two-step lever swing associated with P_i_ release and ADP release. The myo1c ADP•P_i_ state is represented by the crystal structure of a myo1c vanadate co-complex (PDB: 4BYF; (28)); this structure is docked to actin by aligning the lower 50 kDa subdomain with our AM structure. (*B – C*) Putative motion of myo1c and actin during the power stroke when the myo1c tail is anchored to a membrane, as inferred from the structures in (*A*). (*B*) If the actin filament orientation is held fixed with respect to the membrane, the power stroke translocates the actin filament parallel to the membrane. (*C*) If the myo1c tail orientation is held fixed with respect to the membrane, the power stroke rotates the filament, swiveling the barbed end towards the membrane.

Using the crystal structure of myo1c in the presence of Mg^2^ ^+^·ADP·VO_4_ (PDB: 4BYF; (28)) to represent the M.ADP.P_i_ state, we docked FL-myo1c on actin using the lower 50-kDa, that includes loops-2 and -3, and monitored the lever position of the M.ADP.Pi, AM.ADP^A^, AM.ADP^B^, to AM states (Fig. 5). In the pre-power-stroke state, the lever is nearly parallel to the long axis of the actin helix. As the motor progresses from AM.ADP.Pi to AM.ADP^A^ the lever tilts ∼50 degrees in the plane of the actin and then a further ∼20° to the AM state, resulting in a lever position that is nearly perpendicular to the actin filament (Fig. 5). This lever arm angle is very different from other characterized myosins, and it may result in differences in how the motor domain responds to forces aligned with the long axis of the actin filament. Additionally, it has implications for motility relative to the cell membrane (see below).

## DISCUSSION

In summary, we discovered that myo1c binds to actin in a unique orientation that produces a ‘skewed’ power stroke, and that this skewing is further amplified by unique structural features of the myo1c NTE. Together, these features may explain the motor’s ability to turn actin filaments in gliding assays (12, 13). We also found that the conformational changes that myo1c undergoes to release ADP are substantially different from myo1b, and the NTE has a unique structure. These differences may explain differences in force sensitivity. Finally, our new structures allow us to model the working stroke of the full length myo1c molecule, providing insights into function of the native molecule.

### Lever tilting and Force Dependence of ADP Release

To initiate the myo1c lever swing from AM.ADP^A^ to AM.ADP^B^, the L5–lever bridge that connects loop-5 to the lever must break via isomerization of the F74 - Y75 side chain pair. The disruption of this mechanical linkage, which does not happen in myo1b, evidently uncouples the lever position from the nucleotide pocket conformation (Fig. 6; Movies S3-S4). Importantly, we propose that breakage of the L5–lever bridge is rate limiting for the biochemically measured ADP release step. Breakage of the L5–lever bridge would not be force sensitive as little lever movement is required (Fig. 6A). Subsequent rotation of the N-terminal subdomain to a position that allows nucleotide exchange occurs after the lever rapidly tilts and is stabilized by NTE docking and formation of the arginine lock. This aspect of the mechanism is supported by our myo1c^ΔN^ rigor structure, whose nucleotide pocket appears suited for stronger nucleotide binding (i.e., formation of the H-bond interaction belt; Fig. 3G) despite a lever swing that is largely complete. Moreover, biochemical studies of the myo1c^ΔN^ mutant revealed that in the ADP state, most of the motor population released nucleotide much more slowly than wild-type, indicating that ADP release was compromised.

**Fig. 6.**
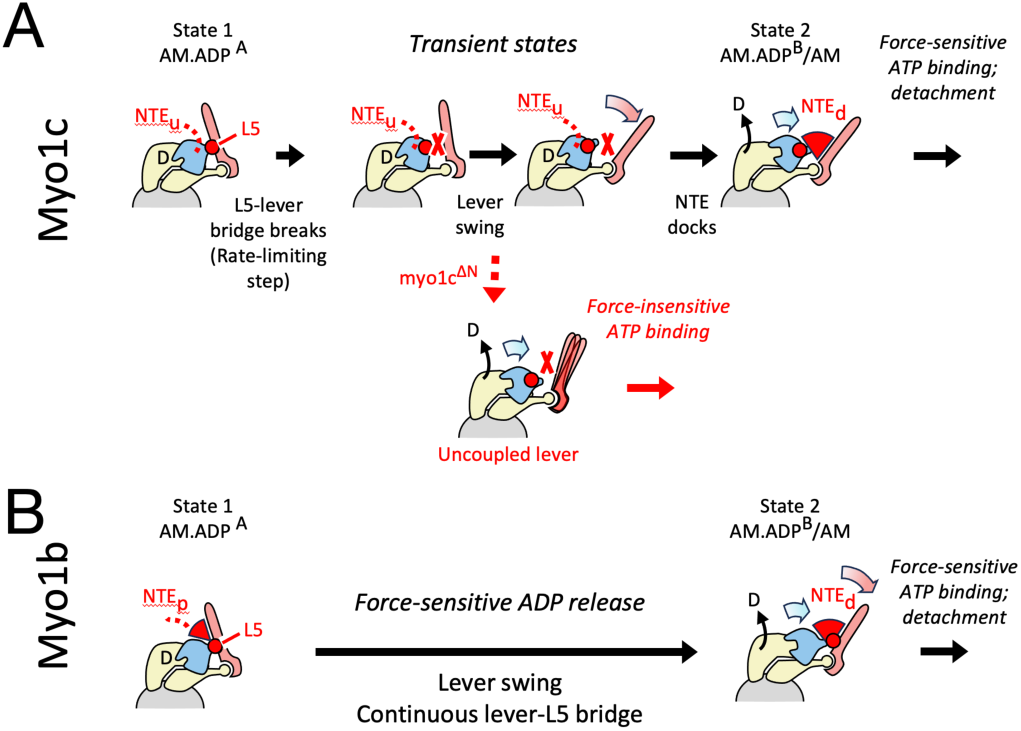
Schematic model of force-mediated nucleotide exchange in actin-bound myo1c and myo1b. (*A*) Force-insensitive ADP release and force-sensitive ATP binding in myo1c. The ADP lever swing is proposed to consist of several force-insensitive substeps: (1) slow detachment of the lever from loop-5 with little lever movement, possibly initiated by isomerization of F77-Y78 in loop-5 (Fig. 4); (2) ADP lever swing, uncoupled from the nucleotide pocket conformation due to plasticity in the loop that connects the converter to SH2 in the motor domain (see Fig. S11); (3) docking of the N-terminal extension, which bridges between the lever and the motor domain and opens the nucleotide pocket slightly to accelerate ADP release (Fig. 3). The undocked or docked NTE is depicted as a dashed red line (labeled ‘N_u_’) or red-colored wedge (’N_d_’), respectively. In the absence of the N-terminal extension (Myo1c^ΔN^; bottom cartoon in panel A), opening of the nucleotide pocket is destabilized, introducing a sub-population that releases ADP slowly. Following ADP release, the docked N-terminal extension tightly couples lever position to nucleotide pocket conformation in the wild-type motor. This confers force-sensitivity to subsequent motor isomerizations that accompany ATP binding, likely involving further movement of the lever that has not been visualized (15). Lacking a docked NTE, ATP binding is uncoupled from lever movement in Myo1c^ΔN^, eliminating force sensitivity. (*B*) Model for force-sensitive ADP release and ATP binding in myo1b. Unlike myo1c, the lever maintains continuous contact with loop-5 in the motor domain during the ADP lever swing, which is proposed to impart force-sensitivity to this step. The myo1b N-terminal extension also has a different role than in myo1c, maintaining interactions with the hydrophobic bridge throughout ADP release and ‘catalyzing’ this step by preferentially stabilizing the transition state (presumed similar to AM.ADP^B^) and final state (AM) over the AM.ADP^A^ state. The red X’s between motor domain and lever (panel A, top row, middle two states) denote severing of the ɸ-bridge. The partially docked myo1b NTE in the ADP^A^ state is depicted as a small red-colored wedge and a dashed red line (labeled ‘N_p_’), while the fully docked myo1b NTE is depicted as a larger red-colored wedge (’N_d_’). Note that AM.ADP^B^ and AM structures are combined as a single depicted image in (A) and (B); this transition is unlikely to contribute to force sensitivity due to minimal motion of the lever in either myo1b or myo1c.

In contrast to myo1c, myo1b maintains its L5–lever bridge throughout the transition from the AM.ADP^A^ to AM states (Fig. 4F–G), which may result in continuous coupling between the N-terminal subdomain and the position of the lever and converter (Fig. 6B; Movie S4). Thus, we propose that the disruption of key bonds that hold ADP in the nucleotide pocket (due to rotation of the N-terminal subdomain) is rate limiting and occurs when the lever has tilted to the AM.ADP^B^ state. Thus, during the AM.ADP^A^ to AM.ADP^B^ lever swing the position of the transition state would be near the AM.ADP^B^ state, resulting in a larger distance parameter and increased force sensitivity of ATP-dependent actin detachment, as measured in the optical trap (10, 11, 38). The transition between AM.ADP^B^ to AM corresponds to an oblique lever movement (14) (Fig. 2F) that is likely to be rapid, and docking of the hydrophobic myo1b NTE is initialized by docking of L10 and L11 in the AM.ADP^A^ state at the beginning of the lever swing.

### Novel compliant element in lever arm linkage

We discovered a compliance in both myo1b and myo1c structures where the SH2 helix connects to the converter. The short linking loop (Myo1c, F627-A628; Myo1b, A632-Y633) found at this junction forms a key mechanical connection between the motor domain and the lever. In other myosin structures (e.g., Myo5 and MYH7) this loop remains mostly rigid during ADP release (Fig. S11), contributing to tight coupling between the nucleotide pocket and the lever swing (39). However, in myo1c, sequence divergence resulting in an alanine (A628) in place of a proline that is highly conserved in non-myosin-I families allows pronounced bending of the SH2-converter linker during the ADP swing (Fig. S11). Nearly identical behavior is observed in myo1b, although this was not previously reported (14). ‘Unhinging’ of this linker in myo1b and myo1c likely facilitates the unique lever positions and tilting angles of myosin-I paralogs.

### Effect of the NTE on ATP binding

Although the transition of myo1c from the AM.ADP^B^ state to AM is accompanied by only a minor (1°) rotation of the N-terminal subdomain, it is accompanied by a slight expansion (or breathing) of the N-terminal subdomain that disrupts the H-bond interaction belt, resulting in a conformation that likely does not tightly bind nucleotide. This finding may explain an interesting effect on ATP-binding kinetics observed for myo1c. Stopped-flow experiments reveal that rigor myosin exists in two states that are in equilibrium; one that binds ATP and another that does not bind ATP, with the non-binding state predominating by more than 3-fold (8, 9). An isomerization in actomyo1c must therefore occur for the ATP to bind; our inability to obtain distinct structures of these two populations from our rigor samples by cryo-EM classification indicates that the structure differences between them must be relatively subtle.

Given its kinetic predominance, our wild-type myo1c cryo-EM AM structure likely represents the conformation that does not bind readily ATP, consistent with our finding of a more open active site. Strikingly, removal of the myo1c NTE results in a biochemical equilibrium of actin-bound myo1c^ΔN^ that favors the state that can bind ATP (16). Indeed, our cryo-EM structure of myo1c^ΔN^ shows a reorientation of the N-terminal subdomain that restores the H-bond interaction belt and re-closes the nucleotide pocket, which is a state that we predict will bind nucleotide tightly. By this interpretation, loosening of the active site by the NTE would tend to accelerate ADP release while simultaneously discouraging ATP binding, consistent with biochemical observations.

### Origins of force sensitivity in the ATP binding step

ATP binding by myosin-I’s is divided into two steps: an initial isomerization in the AM state that permits formation of a weak-ATP binding complex, and a second isomerization to a strong ATP-binding state. As already discussed, the first of these isomerizations likely involves a minor ‘breathing’ movement of the nucleotide active site that enables ATP to interact with the P-loop (Fig. 3F), minimal lever movement (Fig. 1B-C), and is not associated with force sensitivity. The second isomerization, which is linked to force sensitivity by optical trapping and biochemical data (8), has not been characterized structurally but likely involves a ‘post-rigor’ conformation captured in X-ray studies of several other myosins (31, 40, 41). In the post-rigor conformation, the nucleotide pocket closes to interact tightly with ATP, accompanied by movement of the upper-50 kDa domain that weakens the actin interface in preparation for ATP-induced detachment.

To investigate how force sensitivity may be linked with the second ATP-isomerization, we compared our AM myo1c cryo-EM structures with an AlphaFold-generated model of post-rigor ATP-bound myo1c (Uniprot Q5ZLA6x) (42). The AlphaFold model exhibits an approximately rigor-like lever position with a docked NTE (Movie S3). However, unlike AM myo1c or myo1b structures, the AlphaFold post-rigor model does not show ‘unhinging’ of the SH2-converter linking loop (Fig. S11); thus, the coupling pathway from the lever to the nucleotide pocket partially reverts back to the AM.ADP^A^ arrangement in the Alphafold model. Moreover, the Alphafold model shows little lever angle change compared with AM. This contradicts optical trapping force sensitivity measurements, which seem to require that an additional lever movement accompanies the weak-to-strong ATP isomerization (15, 16).

It therefore seems likely that the AlphaFold post-rigor myo1c model does not fully capture the weak to strong ATP-binding isomerization. We speculate that myo1c nucleotide pocket closure may be accompanied by lever rotation somewhat beyond the rigor-like position observed in the AlphaFold model, dislodging (or otherwise introducing strain in) the docked N-terminal extension. A resisting load that drives the lever from this ‘strong-ATP’ position back towards AM would thus favor NTE docking and disfavor strong ATP binding, explaining why this step is force-sensitive. The same mechanism would also explain the observation that deleting the myo1c NTE greatly accelerates the weak-to-strong ATP binding transition (16), since this deletion would allow the motor to more freely fluctuate to the strong ATP-binding conformation.

### The power stroke of full length myo1c

The ability to model the overall power stroke of full length myo1c is an exciting and revealing outcome of this work (Fig. 5). Like most myosins, the lever tilts toward the barbed end of the actin filament (27); however, myo1c starts in the AM.ADP.Pi state in an orientation nearly parallel to the actin filament and progresses to a point that is just past perpendicular. This tilting trajectory is very different from other myosins, where the lever ends in an orientation tilted more towards the barbed end, which leads to the question of why myosin-I evolved this structural adaptation.

A pleckstrin homology (PH) domain in the myo1c tail domain binds directly to phosphoinositides in the lipid membranes (43, 44), so it is relevant to consider the working stroke in relation to the plane of the membrane (Fig. 5). If myosin binds to actin filaments that are fixed in an orientation parallel to the plane of the membrane, the power stroke results in a 10 nm displacement perpendicular to the actin and membrane, resulting in separation of the actin and membrane (Fig. 5B). Alternatively, if the motor is more rigidly fixed to the membrane, the motor could bring the barbed end of the actin filament closer to the membrane (Fig. 5C). This geometry may effectively position the barbed-end of actin to allow polymerization forces to push the membrane. Indeed, *in vitro* experiments show that myosin-I can synergize with Arp2/3 complex to enhance the pushing forces of myosin-I (45).

Finally, the striking finding that myo1c perches on actin at a different angle from other myosins suggests a mechanism for the asymmetric gliding of actin filaments observed in motility assay (12, 13). Experiments in *Drosophila* suggest a role for the myosin-I motor domain in establishing cell and organ chirality during development (13); however, it remains to be determined if the modulation of chiral activity is due to this torque or to modulation of motor kinetics (46). Interestingly, a low resolution structure of actin-bound *Acanthamoeba* myosin-IB also reveals an altered actin perch (25). Thus, this feature has been conserved, so future cell biological experiments will be required to determine the mechanistic role for this structure.

## Acknowledgements

We thank Dr. Daniel Safer and Rick Wike for assistance with protein expression and purification. We would like to acknowledge Dr. Shenping Wu and the Yale Cryo-EM Resource, as well as the Yale Center for Research Computing facility for expert support and maintenance of these facilities. Data collection at Penn was performed at the Electron Microscopy Resource Lab and The Beckman Center for Cryo-Electron Microscopy, University of Pennsylvania (Research Resource Identifier SCR_022375). This work was supported by NIH Grants R01 GM110530 to CVS and R37 GM057247 to EMO, and the National Science Foundation CMMI Grant 15-48571 to EMO. Finally, we would like to thank the reviewers for insightful and helpful comments, which led to new insights and significantly improved the manuscript.

## Data availability

Atomic coordinates and corresponding cryo-EM density maps, including the half maps, masks and FSC curves used to estimate spatial resolution have been deposited in the Protein Data Bank (PDB) and Electron Microscopy Data Resource (EMD) under the accession codes 9CFU/EMD-45563 (myo1c AM.ADP^A^ actin complex), 9CFW/EMD-45565 (myo1c AM.ADP^B^ actin complex), 9CFX/EMD-45566 (wild-type myo1c rigor actin complex, or AM), and 9CFV/EMD-45564 (myo1c^ΔN^ rigor actin complex).

## Code availability

A python script was written for UCSF ChimeraX to perform lever swing analyses and visualizations in Fig. 2, Supplementary Fig. 7, Supplementary Tables S5 and S6. The script is publicly available on gitlab (https://gitlab.com/cvsindelar/lever-swing).

## METHODS

### Protein Preparation

The short, mouse, splice isoform of Myo1C containing the amino-terminus (^1^MESALT…) through all three IQ domains (residues 1-767) immediately followed by the sequence G<u>GLNDIFEAQKIEWHE</u>AA<u>DYKDDDDK</u> that includes a BirA biotinylation site (AviTag; GLNDIFEAQKIEWHE) and a FLAG epitope tag (DYKDDDDK) for purification, was expressed using the Sf9-baculovirus system (43). The myo1c^ΔN^ expression construct was identical to myo1c, except for the removal of the NTE (E2-V10) as described (16). Myosin was purified by FLAG-affinity and ion exchange as previously described(8, 16, 47). Rabbit skeletal muscle actin was purified as previously described (48). Actin polymerization was induced by adding 0.1 volume of 10x KMEI buffer (500 mM KCl, 20 mM MgCl_2_, 10 mM EGTA, 100 mM imidazole, 20 mM ATP, 2 mM DTT) to ∼5 μΜ of G-actin at room temperature for 1 hour. Actin filaments were stabilized using phalloidin at 4 C overnight. Actin filaments were collected by ultracentrifugation (Beckmann Rotors, TLA 120.1, 70000 x g, 30 minutes at 4 C) and the pellet was resuspended in F-actin buffer (10 mM HEPES pH 7.5, 100 mM KCl, 2 mM MgCl_2_, 1 mM DTT). Then, F-actin was incubated with excess myo1c for 1 hour on ice. Actomyosin filaments were collected as a pellet by ultracentrifugation (Beckmann Rotors, TLA 120.1, 70000 x g, 1 hour at 4 C). The pellet was resuspended in F-actin buffer (10 mM HEPES pH 7.5, 100 mM KCl, 2 mM MgCl_2_, 1 mM DTT) at a concentration of 2-5 mg/mL before grid preparation. The myo1c-ADP samples were prepared by dissolving the pellets in F-actin buffer containing 1 mM K_2_ADP.

### Sample freezing and data collection

Samples were frozen on Quantifoil 1.2/1.3 300-mesh (Holey carbon) copper grids; to enhance filament decoration, grids were not glow-discharged but were blotted specially to improve ice quality (49). A sample of 3.0 μL was applied onto the carbon side of the grid using FEI Vitrobot^TM^ Mark IV at 4°C and 100% humidity. The samples were incubated on the grid for 50 s and the extra solution was blotted using two Vitrobot filter papers (Ø.55/20 mm, Grade 595, Ted Pella) for 4 s at 0 blot force. The grids were plunged into liquid ethane at ∼180 °C with a wait time of 0.5 s. The vitrified grids were screened for sample homogeneity and ice thickness in a Glacios 200 kV transmission electron microscope equipped with Gatan K2 summit camera. Electron micrographs for image reconstructions were collected using a Titan Krios at 300 kV, with a Gatan image filter with slit width of 20 eV in nanoprobe mode. The ADP data set was collected using the Krios in the Yale cryo-EM facility, while the two other data sets (wild-type and myo1c^ΔN^ rigor) were collected with at the UPenn Singh center Krios; both microscopes were equipped with a cold-field emission gun. A K3 Gatan summit camera in super-resolution mode was used to collect 1 movie per hole (using serialEM data collection software and EPU software at the Yale and Penn facilities, respectively). The target defocus range was between −2.5 μm and −1.2 μm.

The ADP data set was collected at a nominal magnification of 64K (Yale Krios, GIF plus K3), and cryo-EM structure refinement was carried out using a nominal pixel size of 1.386 Å. The wild-type rigor data set (AM) was collected at a nominal magnification of 64K (UPenn Krios, GIF plus K3) and cryo-EM structure refinement was carried out using a nominal pixel size of 1.36 Å. The myo1c^ΔN^ rigor data set was collected at a nominal magnification of 81K (UPenn Krios, GIF plus K3) and cryo-EM structure refinement was carried out using a nominal pixel size of 1.08 Å. All images were recorded in super-resolution mode. Exposure information for the three datasets is as follows. ADP movies: 40 frames, total dose of 52 counts/Å^2^, total of 3800 exposures; WT rigor movies: 45 frames, total dose 50 counts/ Å^2^, total of 3673 exposures; myo1c^ΔN^ rigor movies: 40 frames, total dose of 50 counts/ Å^2^, total of 3704 exposures. Frame exposure duration for the three data sets was 0.065 - 0.13 sec/frame.

All datasets were processed entirely using CryoSPARC v3 (50). Micrographs were subjected to motion correction (Patch Motion Correction) and CTF estimation before particle picking. Filaments were selected using the cryosparc ‘Filament Tracer’ function. After extraction, particle stacks were subjected to 2-D classification to remove non-filamentous particles. The resulting particles (3088218, 2667431, and 2548251 total particles for ADP, WT rigor, and myo1c^ΔN^ rigor datasets, respectively) were then subjected to helical refinement, followed by optical parameter refinement including defocus and magnification anisotropy. Next, single-particle refinement was performed with a particle-subtracted image stack using a focusing mask comprising the central ∼5 actomyosin subunits. Finally, local refinement was performed using a focusing mask comprising the central myosin subunit plus its neighboring actin trimer. Global resolution estimates for the resulting homogeneous reconstructions (actin plus myosin) were ∼2.7 - 2.8 Å for each of the three data sets.

Multiple attempts were made to identify conformational sub-populations. Focused classification with particle subtraction using various target regions, including the lever and N-terminal extension, failed to identify discrete conformational classes. Variability analysis, also using a variety of different focusing masks, was tried on all three data sets but discrete classes were only identified in the ADP data set. The target region for this latter case was a low-resolution mask encompassing the lever arm. Two sequential variability analysis steps were required to obtain well-defined maps for ADP A and ADP B structures (Fig. S4). In the first step, non-occupied myosin sites were sorted out and a relatively pure population (cluster) of 671620 ADP A particles was identified (Fig. S4A - C). Lever density in one of the other identified sub-populations from this first step appeared to reflect a mixture of lever conformations, and was subjected to a second round of variability analysis. This second step yielded a sub-population of 126845 particles with relatively well-defined lever density corresponding to the ADP B structure (Fig. S4D–E). To reduce potential bias due to masking and other artifacts, variability analysis was performed using alignments obtained prior to the final local refinement step (one myosin plus actin-trimer mask), while final 3D reconstructions for ADP structural states were obtained from particle alignments obtained from the final local refinement step (note that this was a global alignment with all 3088218 particles from the ADP dataset).

Following refinement, the resulting volume pixel sizes were adjusted to the most accurate available values. For the ADP data set, a calibrated pixel size of 1.346 Å was used (37), giving an actin repeat spacing of 27.47 Å (ADP A) and 27.37 Å (ADP B). Actin repeat distances were estimated using the ‘measure rotation’ command of UCSF ChimeraX, applied to an identical actin subunit PDB model fit into two different subunit sites in the density maps (51). For the myo1c^ΔN^ rigor structure, final 3D reconstructions were adjusted to a calibrated pixel size of 1.068 Å (38), giving an actin repeat distance of 27.41 Å. For the WT rigor data set, the pixel size was adjusted so that the resulting actin repeat spacing matched that in the ADP reconstruction: a value of 1.332 Å was selected, giving an actin repeat distance of 27.44 Å. Note that this pixel size is significantly smaller than the calibrated size we used for the ADP data set, which was collected with similar instrument settings but a different microscope (Yale vs. UPenn Krios). This discrepancy may therefore be due to differing instrument calibrations for the two microscopes. Alternatively, it is possible for example that the calibrated pixel size (1.346 Å) is accurate for both microscopes, which would mean that that myo1c ADP release triggers a 1% increase in actin spacing. Conclusions of the current work are not affected by this choice of pixel size.

Initial structure models were built using the model-angelo automated tool (52), and further building and refinement was done manually using Isolde (53), COOT (54) and Phenix (55); Isolde was used for the final refinement cycles. Figures were generated using ChimeraX (56).

