## Supplemental Information for "High resolution structures of Myosin-IC reveal a unique actin-binding orientation, ADP release pathway, and power stroke trajectory"

**Movie S1.** Skew of the myo1c ADP lever swing compared with myo1b, illustrating contributions from both a different binding orientation ('perch') on actin, as well as an inherent difference in the lever swing direction of the two motors. Thick transparent cylinders denote the $AM.ADP^A$ and AM swing endpoints, while thin cylinders depict the hinge axis around which the lever rotates, as obtained by the 'measure rotation' command of UCSF ChimeraX (see Fig. S7 for details). A contacting residue pair between the lever base (F634 and Y629 for myo1b and myo1c, respectively) and loop-5 (Y78 and Y75, respectively) are rendered as van der Waals spheres. Rendering and coloring are similar to Figures 2 and S7, but note that the viewing orientation in the movie is rotated 90° counterclockwise compared to the figures.

**Movie S2**. Molecular morph between myo1c and myob AM structures, highlighting differences of the actin binding regions that support 8° motor domain reorientation on the actin surface. Viewing orientation, coloring and depicted components are similar to Fig. 2A. The thick red cylinder depicts the binding reorientation axis defining the motor domain reorientation. Actin-binding loops (green), alpha helices supporting loop 2 and the activation loop (yellow and chocolate), and actin helices involved in the contacts (gray) are rendered as ribbon cartoons; the remainder of the motor domain is depicted as a alpha-carbon worm trace. The actin surface is depicted as a ~15 Å resolution semitransparent gray isosurface. The binding reorientation axis was estimated using UCSF ChimeraX; two copies of a common reference model (myo1b motor domain) were aligned to the two motor domain models, then the rotation and translation transformations between the two references were computed by the 'measure rotation' command; the cylinders generated by ChimeraX depicts the screw axis that relates the two motor domains by rotation around the axis, accompanied by (minimal) translation along the axis.

**Movie S3.** Depiction of the sequence of available myo1c structures spanning ADP release, ATP binding and actin detachment (see Fig. 6A). Key elements involved in the regulation of force sensitivity and nucleotide exchange are highlighted. Our functional interpretation of this sequence is as follows:

**1. Force-insensitive transition from $AM.ADP^A$ to $AM.ADP^B$:** Starting with the tight ADP-binding '$AM.ADP^{A}$' conformation, rotation of the lever by a few degrees separates Y75 in loop-5 from Y629 (in the lever/converter, leading to severing (red 'X') of the ϕ-bridge. This represents a force-insensitive sub-step. Detachment of loop-5 from the lever frees the nucleotide pocket to partially open, but opening is restrained by a 'belt' of interacting residues spanning switch-I (N157), the P-loop (S107 - G108) and A-loop (N53). ADP release thus remains inhibited. Completion of the ADP lever swing to the 'ADP B' state creates a sufficiently large opening between loop-5 and the lever for the N-terminal extension (NTE; magenta) to dock, stabilizing a rigor-like lever position in another force-insensitive sub step.

**2. ADP release:** Starting from '$AM.ADP^B$, new interactions between the NTE and loop-5 stabilize a slightly more open nucleotide pocket conformation, featuring deformation of the nucleotide belt, which stimulates ADP release and by favoring a subtly expanded nucleotide pocket conformation in the AM state. These observations provide a structural basis for the observed slowing of ADP release when the myo1c NTE is deleted ($myo1c^{\Delta N}$ mutant), from 3.9/sec to 0.9/sec. A 10% sub-population of rigor $myo1c^{\Delta N}$ was detected whose ADP release rate remains similar to wild-type; this sub-population may reflect a portion of the myosins, undetectable by our current cryo-EM methods, that achieve the expanded nucleotide pocket conformation despite the absence of the NTE.

**3. First ATP isomerization and weak ATP binding:** Reversal of the preceding 'breathing' movement reverts the nucleotide pocket to an AM.ADP$^{B}$-like conformation. This prepares the nucleotide pocket for ATP binding, but is disfavored by the docked NTE which favors the 'expanded' AM conformation. This proposed function of the myo1c NTE would explain why its deletion accelerates this initial ATP isomerization step from 18/s to 160/s .

**4. Force-sensitive ATP isomerization (strong ATP binding):** Isomerization to a strong ATP-binding state could be captured by a 'post-rigor' conformation produced by Alphafold (Uniprot Q5ZLA6) (1) in which the nucleotide pocket adopts a closed conformation. Movement of the upper 50-kDa domain while holding the lower 50-kDa domain fixed on the actin surface leads to detachment of the upper-50 from the actin surface. However, there is little difference of the Alphafold model lever angle compared with the AM structure. This feature of the model contradicts optical trapping force sensitivity measurements (see main text Discussion), indicating that the Alphafold model does not fully capture conformational changes that occur in the force-sensitive ATP isomerization.

**Movie S4.** Depiction of the sequence of available myo1b structures spanning ADP release (see Fig. 6B). Rendering and coloring are similar to Movie S3.

**Table S1. Myo1c structural state comparisons by RMSD**. Motor domains from our myo1c cryo-EM structure models were aligned by motor domain only (myo1c residues 12-625; excludes lever and NTE) using the 'matchmaker' tool from UCSF ChimeraX; backbone carbon and nitrogen atomic coordinates only were used for alignment and RMSD calculations (i.e., '*matchmaker #1/P:12-625@CA,N,C to #2/P*' input to the ChimeraX command line). RMSD values for all paired atoms from the alignment are reported. Values are in angstroms.

| | $AM.ADP^{A}$ | $AM.ADP^{B}$ | AM | AM ($MYO1C^{\Delta N}$) |
|---|---|---|---|---|
| **$AM.ADP^{A}$** | 0.00 | | | |
| **$AM.ADP^{B}$** | 1.44 | 0.00 | | |
| **AM** | 1.48 | 0.67 | 0.00 | |
| **AM ($MYO1C^{\Delta N}$)** | 1.33 | 0.74 | 0.50 | 0.00 |

**Table S2. Myo1b structural state comparisons by RMSD**. Motor domains from deposited myo1b cryo-EM structure models were aligned by motor domain only (myo1b residues 16-630; excludes lever and NTE) and RMSD comparisons calculated as in Table S1. Values are in angstroms.

| | $AM.ADP^{A}$ | $AM.ADP^{B}$ | AM |
|---|---|---|---|
| **$AM.ADP^{A}$** | 0.00 | | |
| **$AM.ADP^{B}$** | 1.12 | 0.00 | |
| **AM** | 1.65 | 1.22 | 0.00 |

**Table S3. Myo1c/myo1b structural state cross-comparisons by RMSD**. Different states of myo1c and myo1b cryo-EM structure models were aligned by motor domain only (using residues delineated in Tables S1 and S2) and RMSD comparisons calculated as in Table S1-S2. Values are in angstroms.

| | MYO1C $AM.ADP^{A}$ | $AM.ADP^{B}$ | AM | AM $MYO1C^{\Delta N}$ |
|---|---|---|---|---|
| **MYO1B $AM.ADP^{A}$** | 1.46 | 2.32 | 2.22 | 2.19 |
| **$AM.ADP^{B}$** | 1.68 | 2.08 | 1.88 | 1.97 |
| **AM** | 1.95 | 1.97 | 1.69 | 1.81 |

**Table S4. Myo1c/myo1b intermediate subdomain cross-comparisons by RMSD**. Different states of myo1c and myo1b cryo-EM structure models were compared with each other by aligning upper and lower 50 kD domains (myo1c residues 388-453,455-492,500-525,565-579,165-367,380-387,529-557) using the ChimeraX matchmaker tool as in Tables S1-S3. RMSD values reported are between myo1c and myo1b intermediate domain backbone atoms (residues 12-19,25-71,76-105,584-624 and 16-23,28-74,79-108,589-629, respectively). Values are in angstroms.

| | MYO1C $AM.ADP^{A}$ | $AM.ADP^{B}$ | AM | AM $MYO1C^{\Delta N}$ |
|---|---|---|---|---|
| **MYO1B $AM.ADP^{A}$** | 0.91 | 3.59 | 3.67 | 3.41 |
| **$AM.ADP^{B}$** | 1.39 | 1.94 | 2.15 | 1.95 |
| **AM** | 2.13 | 1.48 | 1.46 | 1.44 |

**Table S5. Lever orientation measurements for myo1c actomyosin cryo-EM and x-ray structures**. These data are plotted in Fig. S7C (red and green points). See Fig. S7 for details of the calculations. Values are in degrees.

| | INCLINATION | AZIMUTH | INCLIN. (MYO1B PERCH) | AZIM. (MYO1B PERCH) |
|---|---|---|---|---|
| **4BYF** | 15.5 | 0 | 14.5 | 19.7 |
| **AM.ADP[A]** | 64.3 | 9.8 | 63.6 | 16.8 |
| **AM.ADP[B]** | 84.6 | 23.6 | 84.7 | 29.1 |
| **AM** | 84.7 | 22.5 | 84.8 | 27.9 |
| **RIGOR MYO1C$^{\Delta N}$** | 80.8 | 20.1 | | |

**Table S6. Lever orientation measurements for myo1b**. These data are plotted in Fig. S7C (black points). See Fig. S7 for details of the calculations. Values are in degrees.

| | INCLINATION | AZIMUTH |
|---|---|---|
| **AM.ADP[A]** | 66.7 | 19.0 |
| **AM.ADP[B]** | 87.8 | 19.1 |
| **AM** | 86.3 | 22.2 |

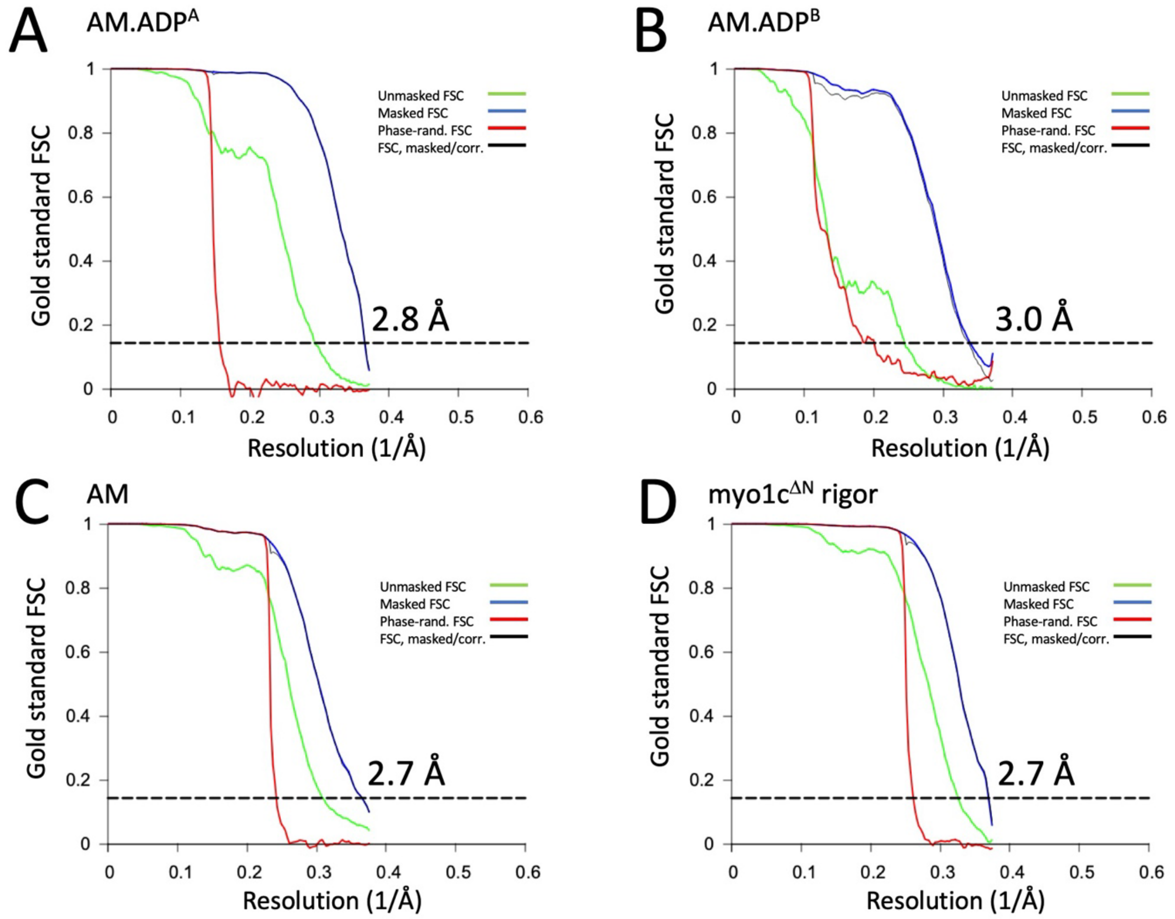


**Fig. S1.** Fourier shell correlation resolution estimates for the four myo1c structures in Fig. 1. Values were computed by Relion (2) (relion_postprocess) using refined half-volume reconstructions and corresponding actomyosin masks generated by cryosparc.

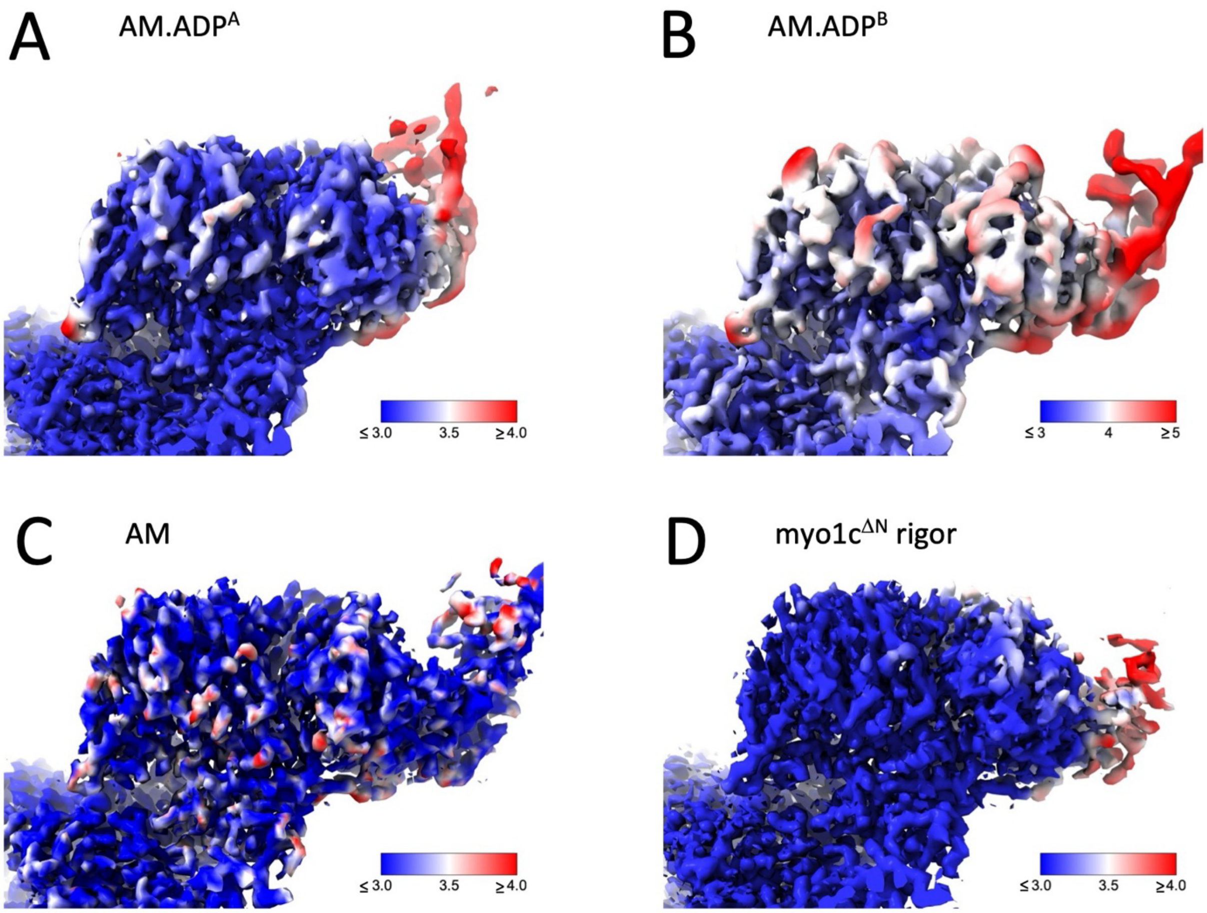


**Fig. S2**. Local resolution estimates of the four myo1c structures in Fig. 1. Local resolution maps were computed using cryosparc ('Local Resolution') (3) from refined half-volume reconstructions and the corresponding actomyosin masks. Here, unlike Fig. 1, the lever is represented with the same isosurface contour level as the rest of the map. Resolution and apparent ordering of the lever arm is significantly greater for the AM state than of any of the other states.

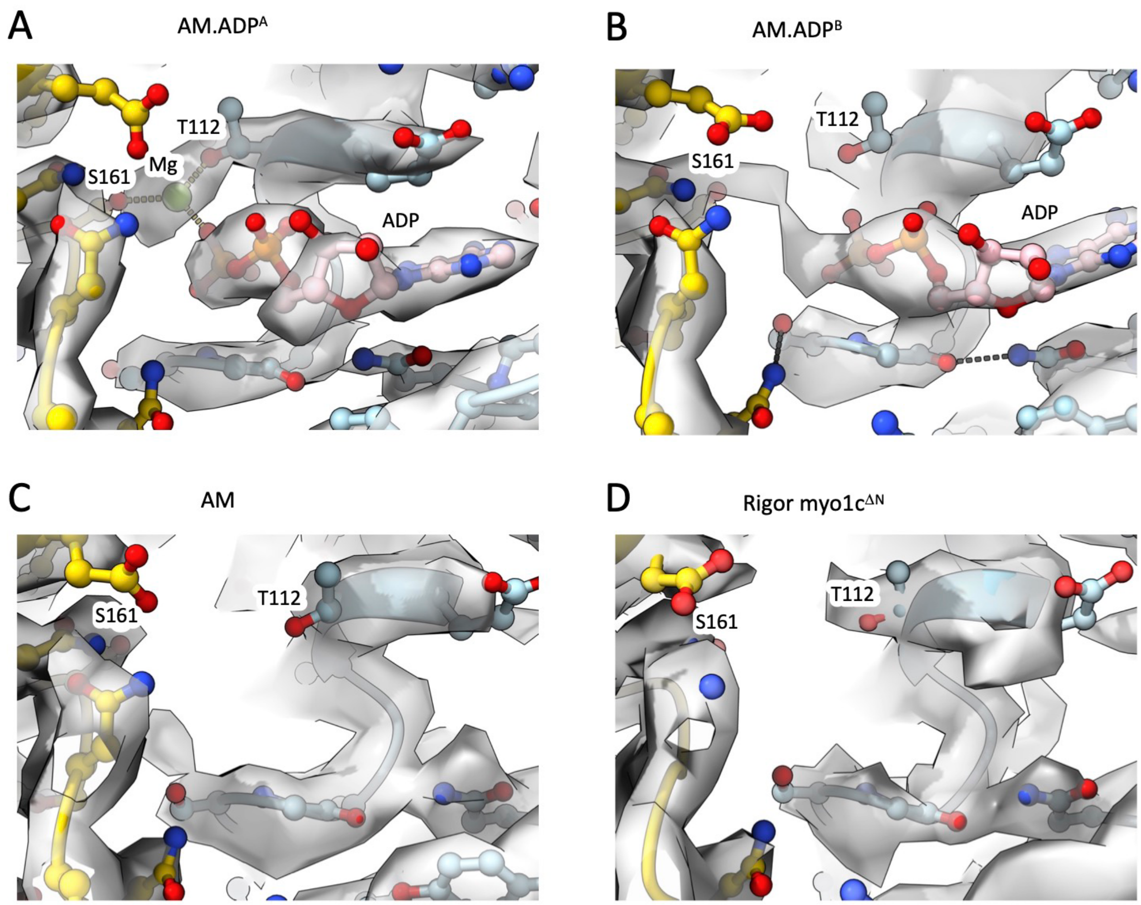


**Fig. S3.** Myo1c magnesium site occupancy is strong only in the AM.ADP$^{A}$ structure. Close-up of the myo1c active site in this figure from a viewing similar orientation as in Figure 1. Cryo-EM map density is rendered as a transparent gray isosurface, with myo1c ribbon diagram superimposed with side chains depicted as balls and sticks.

**A:** Modeled magnesium site (green sphere) is within coordination distance of the ADP beta-phosphate, and T112 and S161 side chains.

**B**: Opening of the active site in myo1c AM.ADP$^{B}$ (see Fig. 3E) perturbs the magnesium site coordination geometry, and localized cation density is not observed. In contrast with myo1c, the myo1b AM.ADP$^{B}$ active site opens less fully, and cation density was still observed (4).

**C,D:** Nucleotide/Mg densities are absent from the rigor structures (AM and rigor myo1c$^{\Delta N}$).

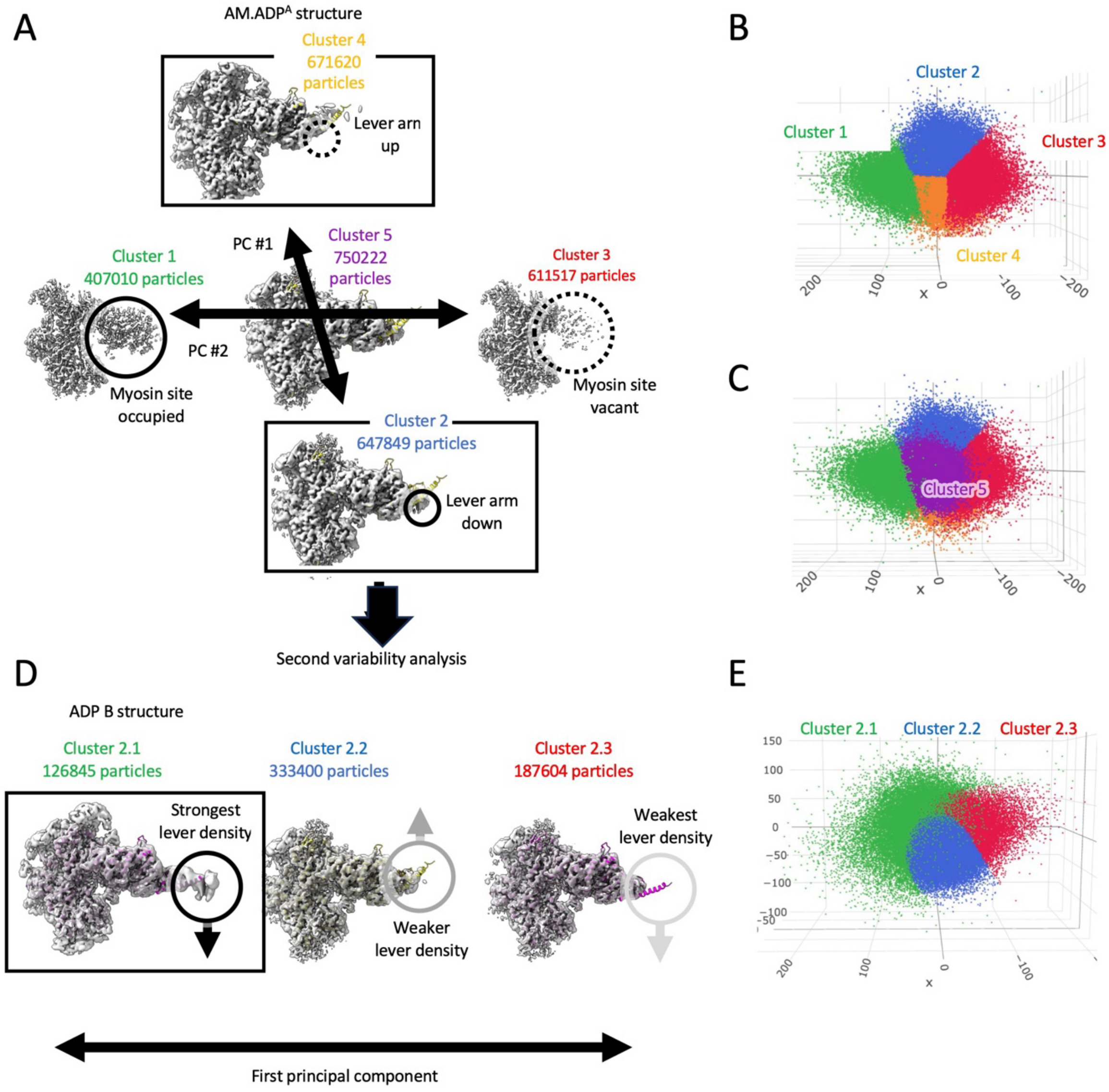


**Fig. S4.** Workflow to sort and reconstruct myo1c 'ADP A' and 'ADP B' particles using variability analysis. A - C, Variability analysis of the full myo1c ADP dataset. After homogeneous high-resolution refinement of the all-particle data set, cryosparc ('Variability analysis') (5) was used to identify principal component axes in the structure latent space (A). The first two principle components were found to correspond to lever arm orientation ('PC #1') and myosin occupancy on the actin filaments ('PC #2'), respectively. Sorting the particles along these principle components using a five-component mixture model (cryosparc 'Variance display') yielded five particle clusters (B-C). The cluster corresponding to the 'up' lever arm position (671620 particles) was assigned to 'ADP A', while the cluster corresponding to the 'down' lever position (647849 particles) indicated a mixture of lever positions and so was subjected to an additional round of variability analysis (D-E). From this second round of variability analysis, a cluster of 126845 particles was identified with a 'down' lever position and designated 'ADP B'.

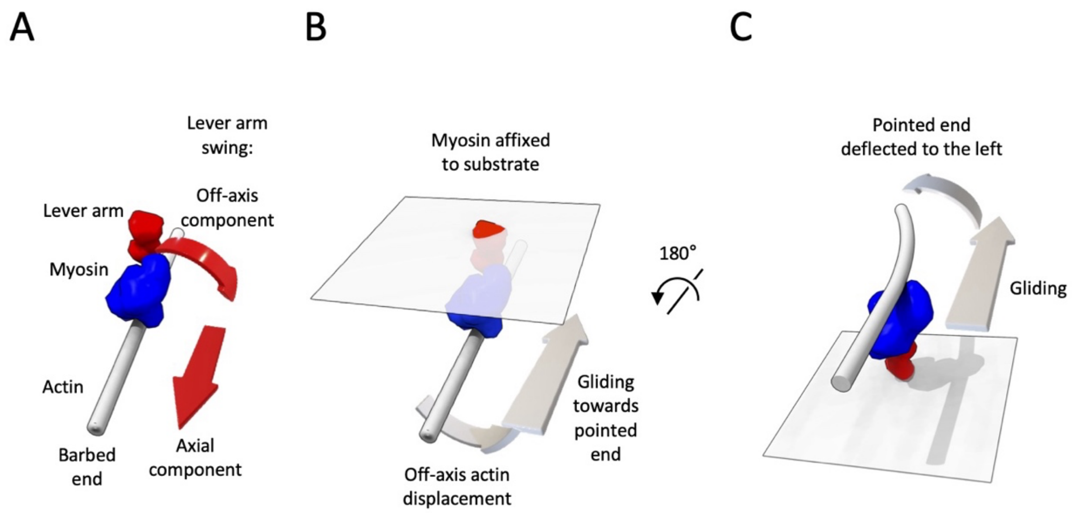


**Fig. S5.** Model for counterclockwise turning bias in myo1c actin filament gliding assays.

**A:** When viewing the actin filament (gray) from its barbed end, the off-axis component of the myo1c lever swing (Fig. 2A-D) moves the end of the lever (red) rightwards as it swings toward the barbed end.

**B:** When the myo1c lever is attached to a fixed substrate, as in a gliding assay, the actin filament moves rightwards with respect to the substrate.

**C:** When the view in B is flipped to place the myosin-coated side of the substrate on top, the myo1c power stroke tries to move the filament to the left. Thus the leading end of a filament would be deflected leftward, potentially leading to counterclockwise circular gliding.

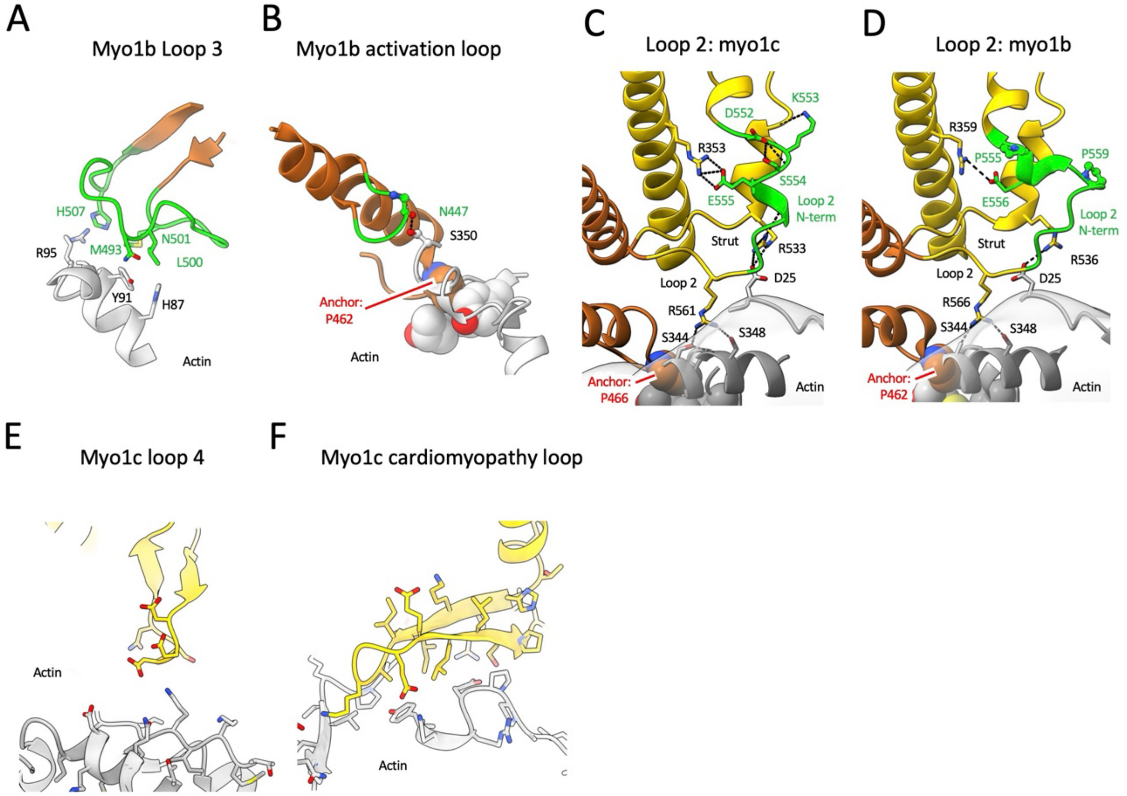


**Fig. S6.** Comparison of myosin 1b lever arm swing and actin interfaces to those of myo1c shown in Fig. 2.

**A,B:** Actin interactions of the myo1b loop-3 and activation loop, with the same viewing orientations as Figs. 2C and 2D, respectively.

**C,D:** Loop-2 structure details and actin interactions in myo1c and myo1b (respectively) that support relative movement of the upper and lower 50 kDa domains. The viewing orientation is similar to Fig. 2E.

**E**: Loose electrostatic contact of myo1c loop-4 with actin, contrasting with closer hydrophobic actin contacts of this loop seen in myo1b.

**F**: Actin interactions and conformation of the myo1c cardiomyopathy loop, which are very similar to those of myo1b.

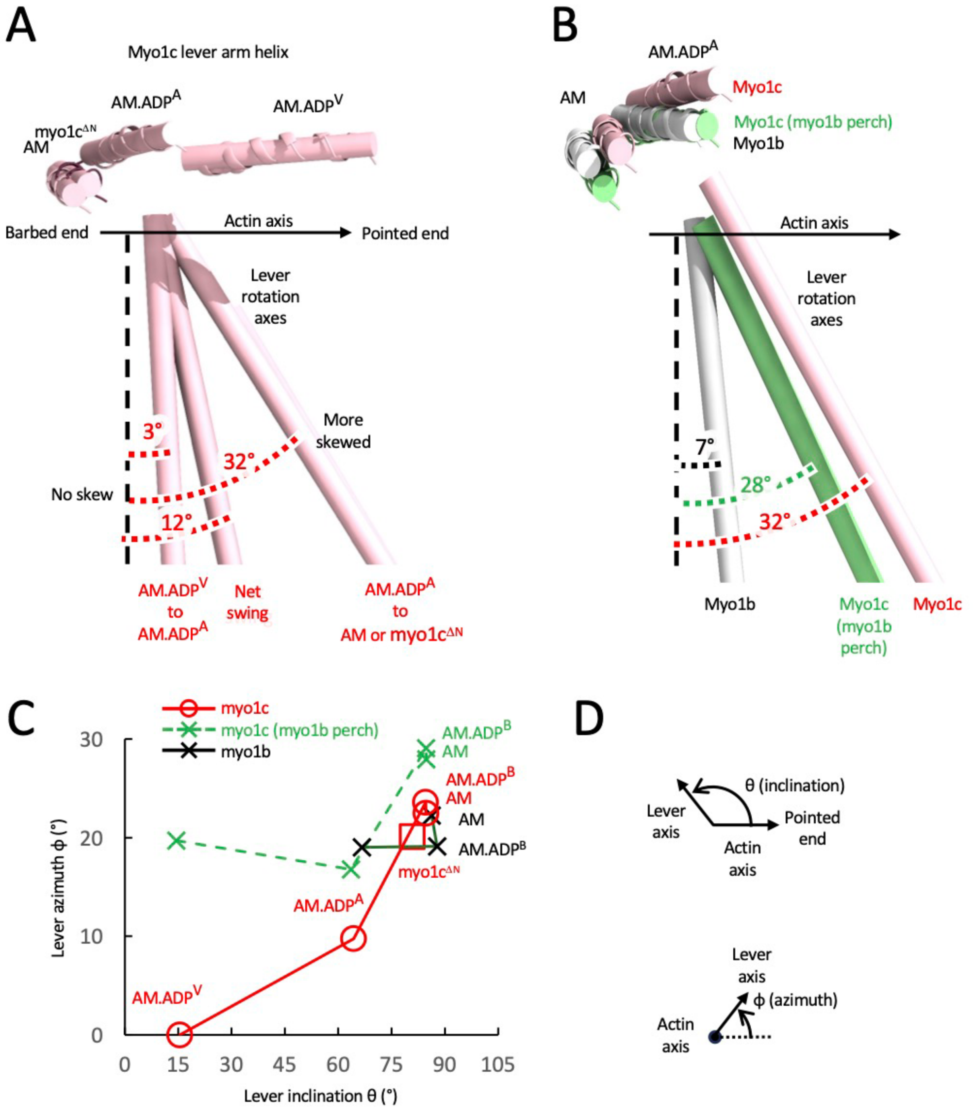


**Fig. S7.** Quantifying off-axis components of myo1c vs myo1b lever arm swings (see also Movie S1).

**A:** Significant off-axis components identified in the modeled myo1c lever swing including the primary power stroke. The top row of pink cylinders, juxtaposed with the myo1c lever helix, depict the lever orientation of modeled AM.ADP$^V$, AM.ADP$^A$ and AM actomyosin models, aligned with respect to the actin filament as in Fig. 2. The lower cylinders depict the axes of rotation determined for each predicted lever transition, as in Fig. 2B. For reference, the black vertical dashed line denotes a lever rotation axis with no off-axis component, as in Fig. 2B.

The pre-power stroke state was modeled by aligning the AM.ADP$^V$ myo1c X-ray structure (PDB: 4byf) (6) by its lower 50-kDA domain (represented by amino acid residues 388-492,500-525,565-579) to that of our myo1c AMD.ADP$^A$ cryo-EM structure, under the assumption that the lower 50-kDa domain in these biochemical states is similarly positioned with respect to actin (7).

The primary lever swing (AM.ADP$^V$ to AM.ADP$^A$) is predicted to have much less off-axis component (3° skewed from a fully parallel swing), than the ADP lever swing (AM.ADP$^A$ to AM; 32° off-axis skew). The overall swing from pre-power stroke to AM models is a composite of the primary and ADP swings, and therefore has a significant off-axis component (12° skew) that falls between those of the sub-swings. Note that the net skew is closer to that of the primary lever swing because the primary lever swing contributes more to the power stroke (primary swing inclination change is ~50° vs. 24° for the ADP swing; Fig. S7C).

All lever rotation axes were estimated using UCSF ChimeraX; different copies of a common lever reference model (4byf amino acid residues 390-410) were aligned to the two lever models being compared, then the rotation and translation transformations between the two lever references were computed by the 'measure rotation' command; cylinders generated by ChimeraX (lever rotation axes) depict the screw axis that relates the two lever models by rotation around the axis, accompanied by translation along the axis. Lever swing off-axis skew angles were calculated using the inner (dot) product of the lever rotation axis and filament axis unit vectors:

$$\theta_{\text{skew}} = 90 - \cos^{-1}(\hat{r} \cdot \hat{z})$$

where $\hat{r}$ is the unit vector along the estimated axis of lever arm rotation, and $\hat{z}$ is the unit vector in the direction of the actin filament helical axis. These unit vectors were computed in UCSF ChimeraX using custom python scripts (lever_axis.py, functions 'lever_axis' and 'helical_axis'; see 'Code availability').

**B:** Myo1c ADP to AM lever swing has a larger off-axis skew angle than myo1b. ADP lever swing axes are shown for myo1b (white), myo1c (pink), and myo1c re-oriented to the myo1b perch orientation (green).

***C:*** Lever angular positions of myo1b and myo1c with respect to actin, represented as a combination of components parallel ('inclination') and perpendicular ('azimuthal') to the actin helical axis. Angle definitions are depicted in Fig. S7D.

Measurements were performed with UCSF ChimeraX (8), using the Angles/Axes/Centroids function ('Axes'), facilitated by a custom python script (lever_axis.py, implementing ChimeraX commands 'helical_axis', 'lever_axis' and 'lever_swing'; see 'Code availability'). The lever orientation was defined using residues 690-710 and 685-705 of the myo1c (current work) and myo1b (PDB 6c1d, 6c1g, 6c1h (4)) structures, respectively. For these measurements, actin filaments were aligned to the myo1c AMD.ADP$^A$ structure using the UCSF ChimeraX 'matchmaker' function, specifying the three actin subunit chains spanning the actin interface of the central myosin subunit (command line equivalent is, i.e. ' match #2/B-D to #1/B-D pairing ss').

Lever inclination ($\theta$) was estimated using the inner (dot) product of the lever orientation ($\hat{l}$) and filament axis ($\hat{z}$) unit vectors: $\theta = \cos^{-1}(\hat{l} \cdot \hat{z})$. The lever azimuth angle ($\phi$) was estimated by taking the projection of the lever orientation unit vector onto the *xy* plane orthogonal to the filament axis (giving an azimuthal orientation vector), and computing the angle between the resulting vector and a reference vector (here taken to be the lever azimuthal orientation vector of the pre-stroke model '4byf').

To approximate the myo1c lever swing if the motor were bound to actin in the myo1b 'perch' orientation (green points; third and fourth columns in Table S4), myo1c coordinates were aligned by their upper and lower 50 kD domains (represented by amino acid residues 388-

492,500-525,565-579 and 165-367,380-387,529-557, respectively) to the myo1b AMD.ADP$^{A}$ filament complex (PDB 6c1d) (4) using the matchmaker function.

**D:** Schematics depicting the definition of lever inclination and azimuthal angles with respect to that actin axis. In this convention, when viewing actin axially from the pointed end a counterclockwise movement of the lever corresponds to a positive azimuthal angle change.

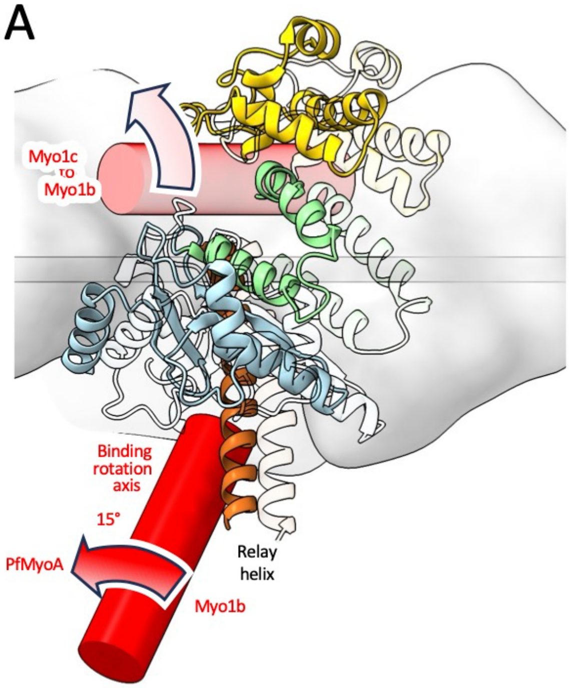


**Fig. S8.** Comparisons of myo1b, myo1c perch orientations vs other myosins.

**A:** Comparison of Plasmodium falciparum myosin (PfMyoA) and myo1b AM (rigor) actomyosin structures reveals a motor binding reorientation axis (red cylinder) roughly parallel to the relay helix. The myosin lever rotation axis also runs roughly parallel to the relay helix (see Fig. 2B). Thus, compared with myo1b, PfMyoA 'rocks' on actin around the same axis as the lever rotation, so that the 'skew' of the lever swing with respect to the filament axis would not be much affected. A similar conclusion is indicated for Myo6, which shares a similar actin binding orientation as PfMyoA (9). This contrasts with the binding orientation difference of myo1c (semitransparent red cylinder, same as the red cylinder in Fig. 2A-B), which leads to significant lever swing skew (Fig. S7). Several other characterized myosins (Myo1a, *Dictyostelium* Myo1e, MYH7; MYH11; Myo15a) have binding orientations on actin similar to myo1b (10-14), indicating their 'perch' also does not strongly skew the lever swing.

**B:** A similar comparison of Myo5 and myo1b AM (rigor) states reveals very similar 'rocking' behavior as PfMyoA and Myo6, although the reorientation magnitude (8°) is less. Thus, while myosins can adopt a range of binding orientations on actin (9), the binding orientation of myo1c is distinct in skewing the lever swing away from the filament axis.

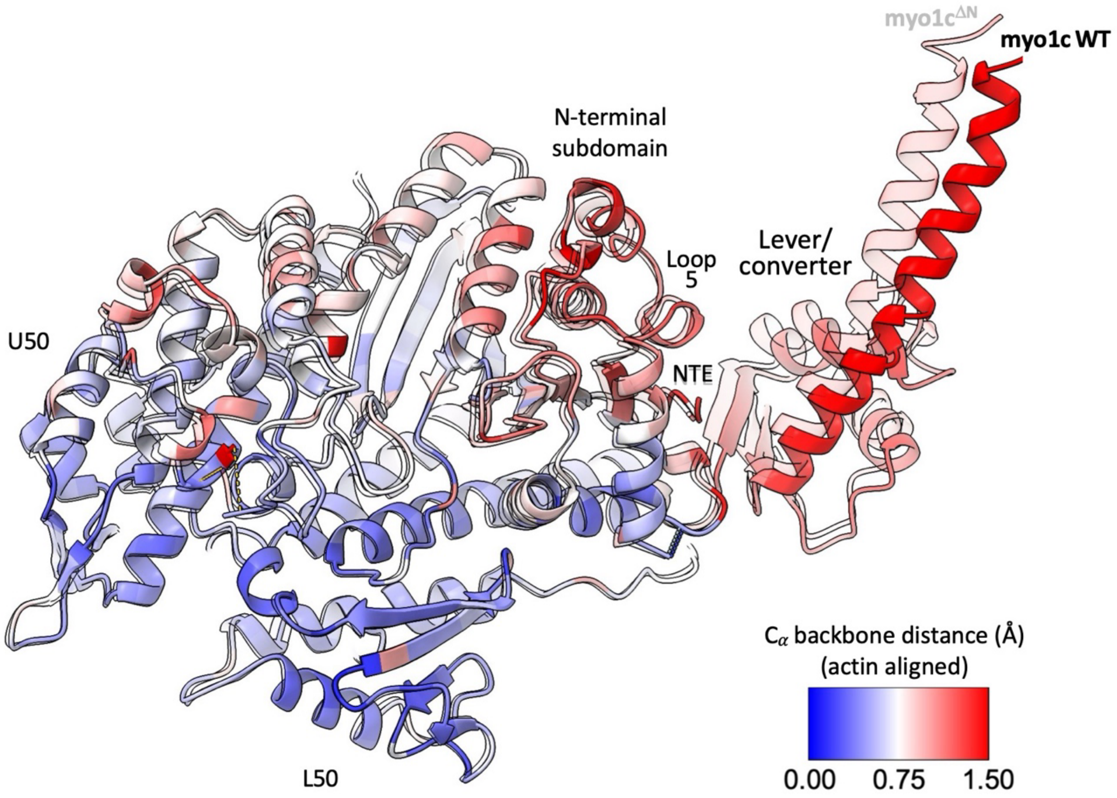


**Fig. S9.** Comparison of the wild-type myo1c (solid ribbons) and myo1c$^{\Delta N}$ (semitransparent ribbon) rigor (AM) cryo-EM structures by backbone distance, revealing concerted 'breathing' movements propagating from the nucleotide pocket to the lever. Structures are aligned by least-squares fitting of actin subunits *i* and *i* + 2 that interface with the motor. The distance color scale ranges from 0 Å (blue) to 1.5 Å (red).

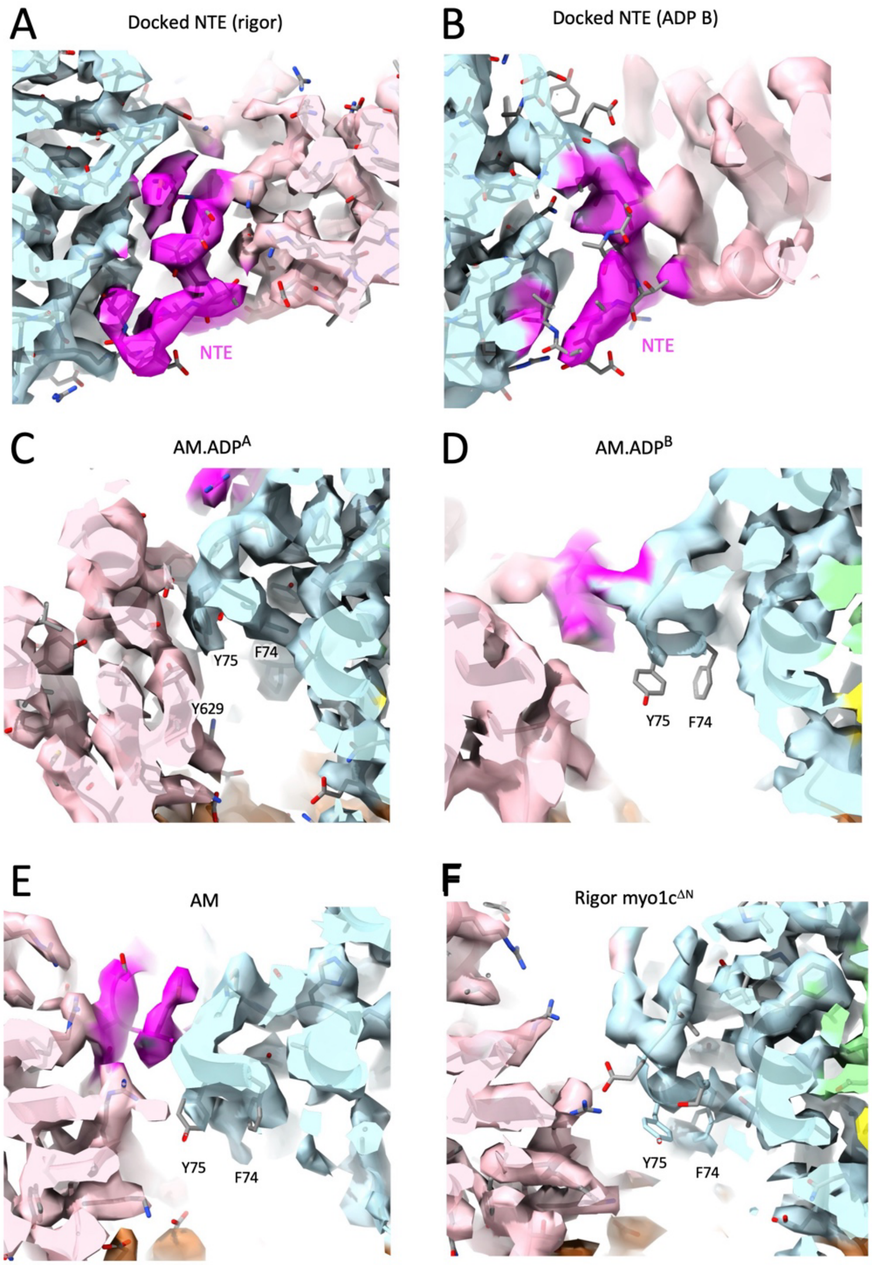


**Fig. S10.** Density features associated with myo1c N-terminal extension docking site shown in Fig. 4. **A–B:** Densities of the docked myo1c AM and AM.ADP$^{B}$ N-terminal extension, respectively; viewing axis is in the plane of Fig. 4C–E, looking from top to bottom. Density is rendered as a semitransparent isosurface, using the same coloring scheme as the molecular rendering in Fig. 4 (intermediate domain, light blue; N-terminal extension, magenta). **C–F:** Isosurface density renderings of the N-terminal extension docking site for all four myo1c structures. View is identical to that of Fig. 4C–E. Hydrophobic bridge side chains F74 and Y75 are well ordered in the AM.ADP$^{A}$ state, but change rotamers and become significantly less well ordered upon dissolution of the bridge in the AM.ADP$^{B}$ and AM structures.

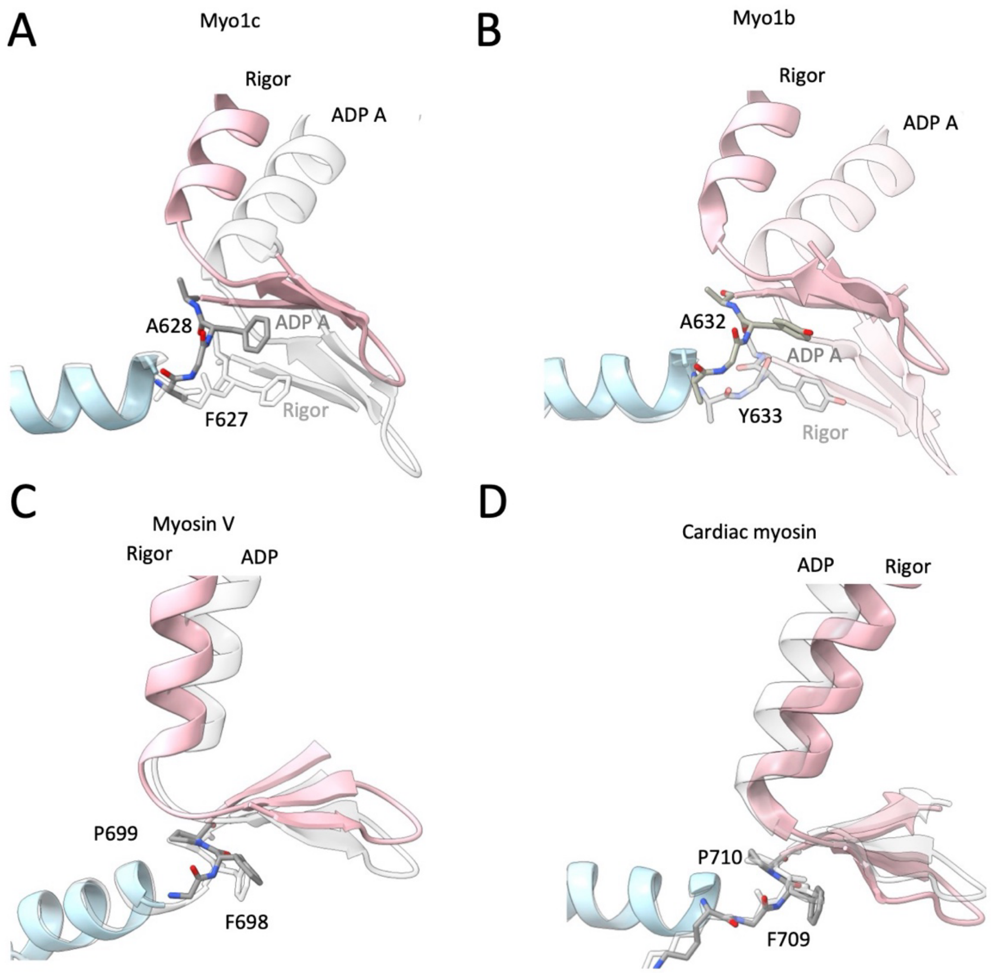


**Fig. S11.** 'Un-hinging' of the SH2 - converter loop in myo1c and myo1b. Viewing orientation is similar to Fig. 1. The SH2 helix (light blue), lever helix and converter beta sheet (pink) are rendered as ribbon cartoons. A two-residue sequence from the SH2-converter linker (FA in myo1b, FY in myo1c, PF in myo5 and cardiac myosin) is rendered as a stick diagram.

**A, B:** In myo1c and myo1b, A628 and A632 (respectively) introduce flexibility in the SH2-converter linkage, permitting the lever to move > 20° during ADP release.

**C, D:** In other myosins including MyoV (PDB 7PM6 and 7PLU) (12) cardiac myosin (PDB 8EFD, 8EFE) (13), the alanine in the converter linkage is substituted with a proline. This substitution restricts mobility in the SH2-converter linkage, likely accounting for reduced lever movements that accompany ADP release in these other motors.
